# Non-canonical Metabolic and Molecular Effects of Calorie Restriction Are Revealed by Varying Temporal Conditions

**DOI:** 10.1101/2024.02.15.580502

**Authors:** Heidi H. Pak, Allison N. Grossberg, Rachel R. Sanderfoot, Reji Babygirija, Cara L. Green, Mikaela Koller, Monika Dzieciatkowska, Daniel A. Paredes, Dudley W. Lamming

## Abstract

Calorie restriction (CR) extends lifespan and healthspan in diverse species. However, comparing *ad libitum* (AL) and CR-fed mice is challenging due to their significantly different feeding patterns. CR-fed mice consume their daily meal in approximately 2 hours, subjecting themselves to a prolonged self-imposed fast each day. To gain deeper insights into the effects of CR, we conducted a comprehensive examination of how AL and CR-fed mice respond to tests performed at various times relative to the completion of their once-daily CR meal. Our findings reveal that many well-known effects of CR, including its impact on insulin sensitivity, result from the specific temporal conditions. CR animals exhibit a divergent response to insulin, and this response varies based on the time elapsed since the CR-fed mice consumed their food. Utilizing an unbiased metabolomics approach, we discovered that the effects of CR on circulating metabolites are heavily dependent upon the time-of-day and feeding regimen. Finally, while it is widely believed that CR functions in part by reducing activity of the kinase mTORC1, our study suggests that the observed differences in mTORC1 activity between AL and CR-fed mice are dependent upon both fasting duration and the specific tissue examined. Furthermore, we find that the metabolic effects of CR are independent of hepatic mTORC1. Our results shed new light on the physiological, metabolic, and molecular effects of a CR diet, and highlight that much of our understanding of the effects of CR are related to when, relative to feeding, we choose to examine the mice.

## Introduction

Calorie restriction (CR) is the gold standard for geroprotective interventions ^1^. However, the widespread difficulty in adhering to a CR diet has sparked significant interest in uncovering the physiological and molecular mechanisms that mediate the beneficial effects of CR. While many different molecular pathways have been proposed to mediate the effects of CR on health and longevity, a comprehensive understanding of how CR confers its benefits remains elusive. A potentially complicating factor in our quest to understand the mechanism for CR is that CR- and AL-fed mice have dramatically different eating patterns. In a CR regimen, animals consume their entire daily allotment of food within approximately ∼2 hours, leading to a prolonged involuntary fast between meals^2,3^.

We recently demonstrated that this daily prolonged fast between meals is necessary for the metabolic and geroprotective effects of a CR diet, and the imposition of a prolonged daily fast alone without decreased calorie consumption is sufficient to recapitulate many of the metabolic and molecular effects of a CR diet ^3^. While conducting these studies, we recognized that the extended time between meals for CR mice poses challenges for interpreting physiological and molecular differences between CR- and AL-fed mice for two reasons. First, determining the appropriate time to assess the molecular and physiological effects of CR relative to *ad libitum* (AL) fed controls is challenging due to the sharply different eating patterns of CR and Al mice. Second, CR mice may have adapted to daily fasts, while AL-fed mice rarely experience fasting. Consequently, when comparing AL and CR-fed mice, we are contrasting mice that have adapted to fasting with those that have not^3,4^.

As but one example, when examining the response to a glucose or insulin challenge, researchers routinely fast animals for a period between 6-16 hrs to minimize variability and bias in blood glucose measurements ^5–7^. This approach not only assesses the animal’s response to the administration of glucose or insulin, but also their response to the preceding fast. Furthermore, CR-fed animals, which are subjected to a prolonged fast each day, may have adapted to a fast, whereas *ad libitum* fed mice, are rarely subjected to fasting. Thus far, no study has comprehensively examined how the phenotypes of AL and CR-fed mice evolve as the time since their most recent meal increases, with most studies comparing AL and CR-fed mice at a single timepoint.

Here, we investigated the physiological and molecular responses of mice fed an AL or CR diets, varying the time since their most recent meal from 4 to 24 hours. Notably, we find that while CR-fed mice exhibited an overall improved response to glucose, an improved response to insulin relative to AL-fed controls was only evident at specific timepoints following their last meal. Specifically, the response to insulin, assessed via intraperitoneal (IP) administration, displayed significant variations in CR-fed mice based on the length of the preceding fast, a pattern not observed in AL-fed mice. Additionally, our research revealed time-of-day-dependent effects on insulin levels in CR-fed mice, highlighting the crucial role of insulin in suppressing hepatic glucose production during an extended fast. Lastly, contrary to the prevailing theory that CR promotes health and longevity by reducing mTORC1 activity^8^, our findings indicate that hepatic mTORC1 activity is strongly regulated by the feeding-fasting status, and is not constitutively suppressed by CR. Conversely, in muscle, mTORC1 activity is elevated in a time-dependent manner under a CR feeding regimen. Our study underscores the significance of considering temporal conditions in comprehending the physiological, metabolic, and molecular effects of CR, and underscores the necessity to consider these factors when evaluating the effects of dietary interventions.

## Results

### Fasting duration produces differential response to exogenous insulin in mice fed a CR diet

We initiated a study with 9-week-old male C57BL/6J mice, dividing them into two groups: AL (never subjected to fasting) or 30% CR (exposed to daily fasting). Both groups were fed Envigo Teklad Global 2018; a diet that extends lifespan and improves metabolic health in CR-fed mice^3,9^. The CR mice were fed once per day at the beginning of the light cycle (Zeitgeber Time, ZT 0) a common practice in CR studies, including the National Institute on Aging (NIA) aged rodent colony^9–12^. We measured the bodyweight for the efficacy of the diet, and as expected, male mice on CR had decreased weight gain and decreased adiposity, with maintenance of lean mass (**Figs. 1A-D).**

**Figure 1:**
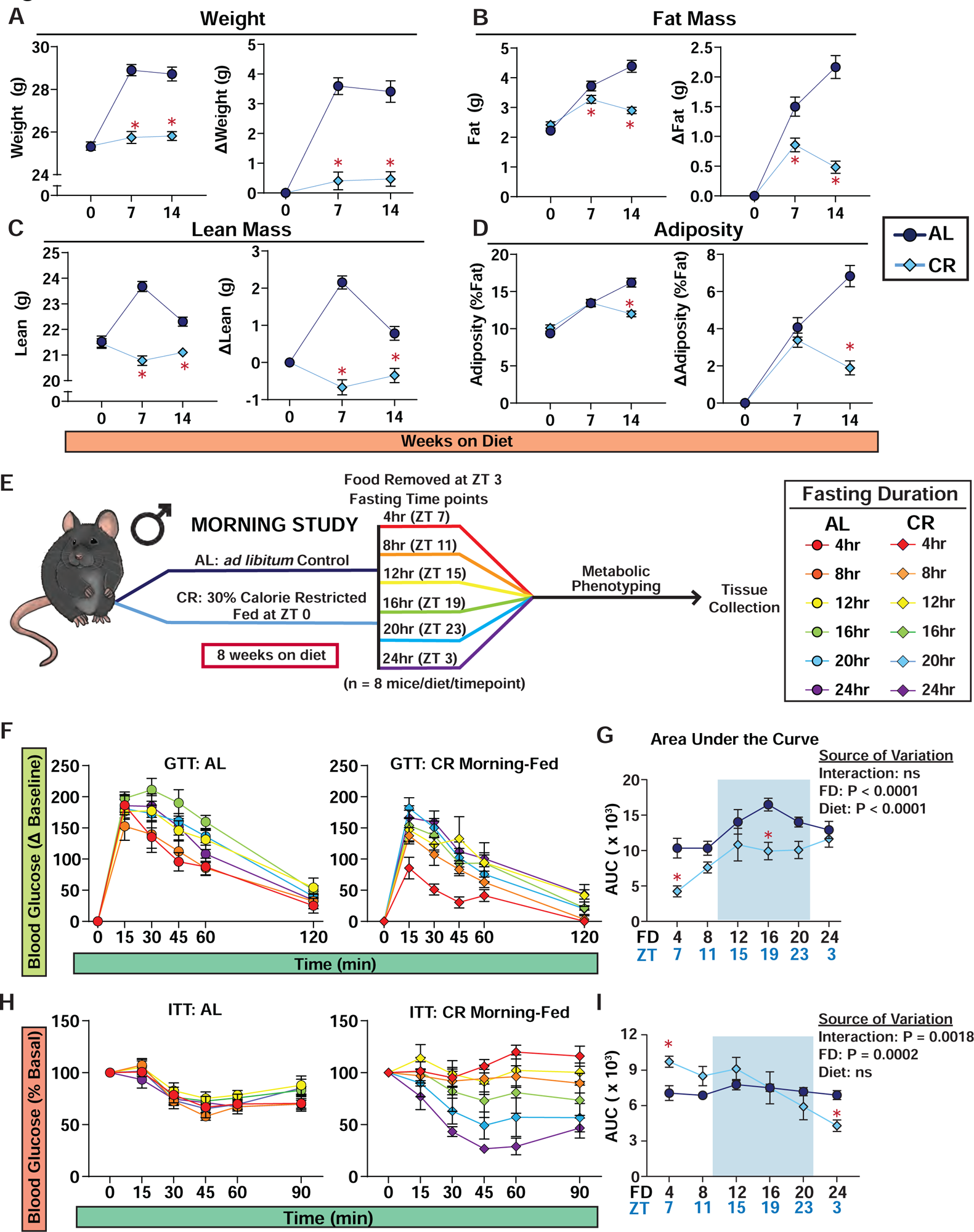
Response to insulin is dependent upon the length of the preceding fast in CR-fed mice. (**A-D**) Body composition measurement of male C57BL/6J mice under Morning-Fed conditions (AL, n = 24 and CR, n = 24 biologically independent mice) *p<0.05 AL-fed vs. CR-fed mice at each time point, Sidak’s test post two-way repeated measures ANOVA. Data represented as mean ± SEM. (**A**) Total body weight with change in body weight from baseline. (**B**) Fat mass with change in fat mass from baseline. (**C**) Lean mass with change in fat mass from baseline. (**D**) Adiposity with change in adiposity from baseline. (**E**) Experimental design of Morning-Fed study. (**F-G**) Glucose tolerance tests of AL and CR-fed C57BL/6J male mice (**F**) and AUC (**G**); food was removed from all groups at ZT 3, and each group was then fasted for varying lengths of time. (**H-I**) Insulin tolerance tests of AL and CR-fed C57BL/6J male mice (H) and AUC (I). (**F-I**) *p<0.05 AL-fed vs. CR-fed mice at each time point, Sidak’s test post two-way ANOVA. The overall effect of diet, fasting duration (FD), and the interaction represent the p-value from a two-way ANOVA. Data represented as mean ± SEM. AUC, area under the curve. See **Supplementary Figures 1-2** for comparisons of AL and CR-fed mice at each time point.

To examine the metabolic response to glucose and insulin following fasting for different lengths of time, we conducted glucose and insulin tolerance tests (GTT and ITT, respectively) at various fasting time-points. For each test, both AL and CR-fed mice were given access to food from ZT 0-3, and at the conclusion of this period, all remaining food was removed from all groups. We then randomized the fasted AL and CR-fed mice into 6 groups of 8 mice each, with approximately the same weight and body composition. We then performed the indicated assay (e.g., glucose tolerance test), with the first group examined at ZT 7, and a different group examined every four hours after that (**Fig. 1E**).

During the GTT, we noticed that the fasting blood glucose varied between diet groups and fasting timepoints, contributing to an observed difference between groups (**Supplementary Fig. 1A-C**), prompting us to calculate the delta from the baseline to calculate the AUC (**Supplementary Fig. 1D**). Both AL and CR-fed C57BL/6J males (B6M) had the lowest area under the curve (AUC) after 4 hrs of fasting as compared to other fasting timepoints (**Figs. 1F-G**). While CR-fed mice had a lower AUC than AL-fed mice at all time points, this was only statistically significant after a fast of 4 or 16 hours (**Fig. 1G**).

In contrast, while the response to insulin of AL-fed male mice remained consistent regardless of the length of the fast, the response of CR-fed male mice to insulin was highly dependent on the length of the fasting period (**Figs. 1H-I, Supplementary Fig. 2**). While the literature broadly notes that CR improves insulin sensitivity in mammals ^1^, CR-fed mice fasted for only 4-12 hrs had blood glucose levels that were essentially resistant to insulin administration. Only after 20 or 24 hrs of fasting did CR-fed mice exhibit improved insulin sensitivity relative to AL-fed mice (**Fig. 1H, Supplementary Fig. 2)**.

### Male CR mice tightly maintain glucose homeostasis during a prolonged daily fast

To investigate the unique response to insulin in CR mice, we conducted a meal-stimulated insulin secretion (MSIS) test, measuring insulin levels post feeding rather than after glucose stimulation. This approach aimed to assess insulin levels under natural feeding conditions. Both AL and CR groups had access to food from ZT 0-3. Subsequently, we divided the mice into six groups, subjecting them to fasting periods of either 4, 8, 12, 16, 20 or 24 hrs as in the experiments presented in **Figure 1**. Following the fast, we collected fasting blood and then provided the mice with food for two hours, after which we then collected fed blood (**Fig. 2A**).

**Figure 2.**
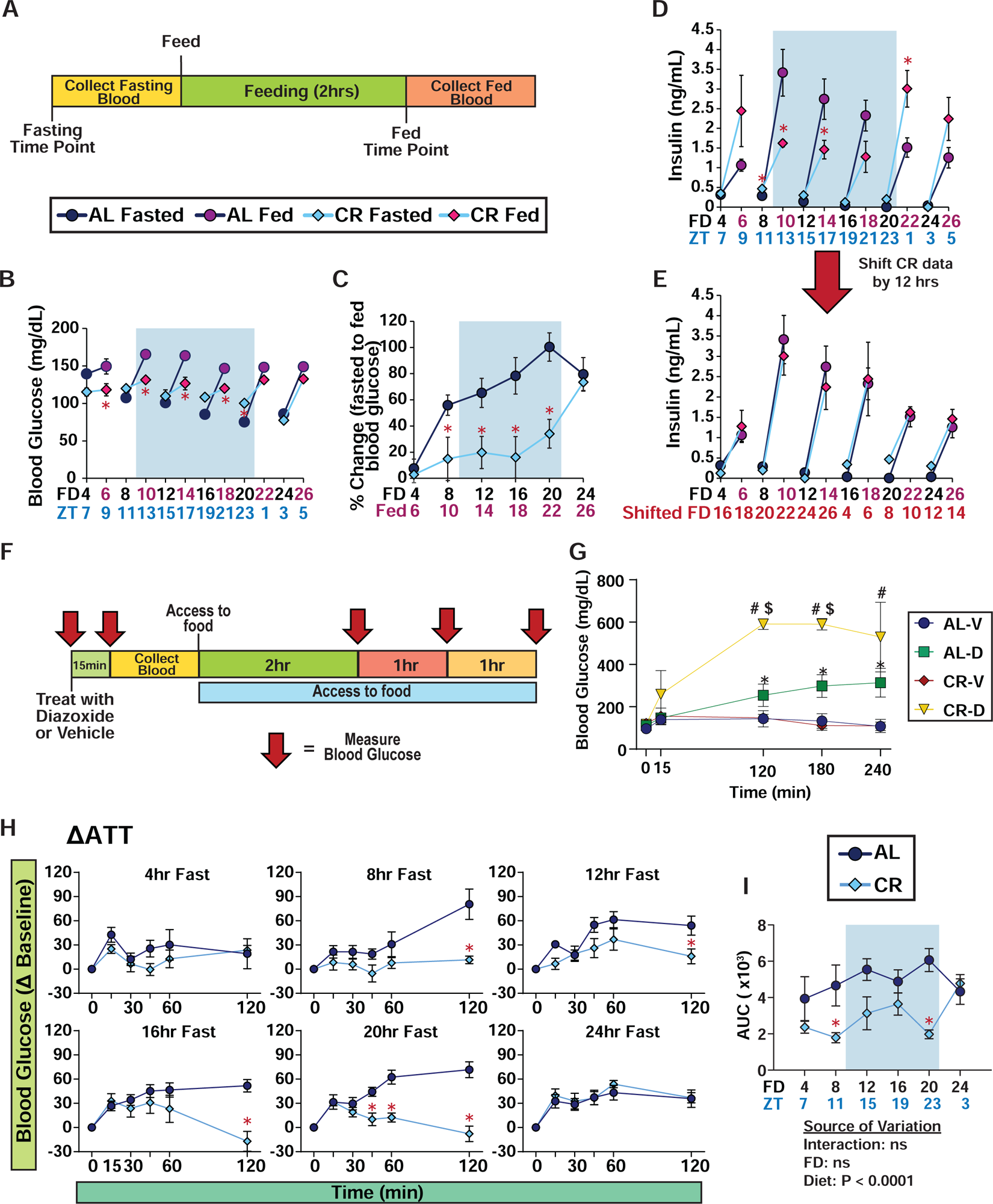
Plasma insulin level after feeding is dependent on time of day, not fasting duration. (**A**) Experimental design of meal stimulated insulin secretion test. (**B-C**) Blood glucose level of mice fasted for varying lengths of time, prior to and subsequent to feeding; FD axis indicates length of time since fasting initiation; red axis label indicates mice refed for 2 hours (**B**), and percent change in blood glucose level from fasted to fed state (**C**). (**A-C**) n=7-8 AL-fed and CR-fed biologically independent mice at each time point. (**D-E**) Plasma insulin level of mice fasted for varying lengths of time, prior to and subsequent to feeding (**D**), and data presented with CR data shifted by 12 hours (**E**). (**D-E**) n=6-8 AL-fed and CR-fed biologically independent mice at each time point. (**F**) Experimental design of acute diazoxide treatment experiment. (**G**) Blood glucose level of mice treated with either vehicle or diazoxide after a 12 hr fast, n=6 biologically independent mice per group (AL-V, AL-D, CR-V and CR-D) (**H-I**) Alanine tolerance test of male C57BL/6J mice after during respective fasting timepoints within each diet (**H**), area under the curve (**I**). (**H-I**) AL, n = 7-8 per time point; CR, n = 7-8 per time point biologically independent mice. (**B-E**) *p<0.05 AL-fed vs. CR-fed mice at each time point, Sidak’s test post two-way ANOVA (G-I) *p<0.05 AL-fed vs. CR-fed mice at each time point, Sidak’s test post two-way ANOVA. The overall effect of diet, fasting duration (FD), and the interaction represent the p-value from a two-way ANOVA. Data represented as mean ± SEM. AUC, area under the curve. See **Supplementary Figure 3** for comparisons of AL and CR-fed mice at each time point.

Intriguingly, CR-fed male mice exhibited resistance to fasting-induced changes in blood glucose. While AL-fed mice displayed ∼46% reduction in fasting blood glucose over 20 hours, CR-fed mice showed only an ∼15% decrease during such a fast (**Fig. 2B**). Following a meal challenge, CR-fed mice were able to maintain a stable blood glucose level, while AL-fed mice exhibited 50-100% increase in blood glucose level following feeding (**Figs. 2B-C**). After 24 hours of fasting – which exceeds the normal length of time between meals for CR-fed mice – AL-fed and CR-fed male mice had almost identical fasting and refed blood glucose levels (**Fig. 2B-C**).

Contrary to our expectations that insulin levels would parallel blood sugar levels, we were surprised to find that the plasma insulin levels of both AL and CR mice were independent of fasting duration and diet (**Fig. 2D**). Although we anticipated lower fasting and refed insulin levels in CR-fed mice compared to AL-fed mice, this was only evident at certain time points. CR-fed mice refed after 4, 20, or 24 hours of fasting had higher refed insulin levels than their AL-fed counterparts. Closer examination revealed that the refed insulin pattern of CR-fed mice appeared to be offset by 12 hours from that of AL-fed mice. When we shifted the CR data by 12 hrs, aligning the data based on the start time of the major eating periods, we observed identical refed insulin patterns in CR and AL-fed mice (**Fig. 2E, simplified graph shown in Supplementary Fig. 3A-B**).

To determine if the stable blood glucose level in CR-fed mice was driven in part by differences in insulin secretion, we tested the impact of inhibiting insulin secretion using diazoxide, a well-established drug that inhibits insulin secretion^13–15^. We performed this test after a 12 hour fast because at this timepoint both AL-fed and CR-fed mice had similar fasting blood glucose levels and insulin response while having different refed insulin levels. Mice were either treated with vehicle or diazoxide, and blood glucose level was measured pre- and post-treatment (**Fig. 2F**). We expected to observe similar level of blood glucose level between AL- and CR-fed mice with the initial administration of diazoxide; however, even without the consumption of food or glucose administration, diazoxide-treated CR-fed mice showed a rapid increase in blood glucose to over 600 mg/dL, whereas the blood glucose level of diazoxide-treated AL-fed mice increased more modestly (**Fig. 2G**). This data suggests that during a fast insulin plays a critical role in inhibiting hepatic glucose output in CR-fed animals.

To confirm that hepatic gluconeogenesis is suppressed in CR-fed mice, we performed an alanine tolerance test (ATT), which measures glucose production from this substrate^16^ (**Fig. 2H-I, Supplementary Figs. 3C**). Supporting our hypothesis, CR-fed mice exhibited reduced glucose production from alanine compared to AL-fed mice at all time points up to 20 hours (**Figs. 2H-I and Supplementary Fig. 6D**). Together, our data suggest that CR mice can maintain tight blood glucose levels during a fast, in part by suppressing endogenous glucose production.

### Response to a challenge is conserved between sexes fed a CR diet

Recognizing sex as an important factor in the response to dietary interventions, including CR ^1,9^, we extended our study to examine the response of female C57BL/6J (B6F) mice to glucose and insulin at various fasting durations (**Supplementary Fig. 4A**). In line with our observations of male mice, CR-fed B6Fs had decreased weight gain, predominately due to decreased accretion of lean mass (**Supplementary Figs. 4B-E).**

CR-fed B6Fs had improved glucose tolerance at all fasting durations compared to their AL-fed counterparts, which was statistically significant after a 12 or 20 hour fast (**Supplementary Fig. 5**). Additionally, the response to insulin in CR-fed B6F mice paralleled that of males, with CR-fed females fasted for 4-12 hrs displaying resistance to insulin, both in absolute terms and relative to AL-fed mice, with CR-fed mice showing improved insulin sensitivity only after 20 hours of fasting (**Supplementary Figs. 6**).

When conducting the MSIS test in CR-fed females, we once again observed tight blood glucose control (**Supplementary Figs. 7A-B**). As we observed in males, CR-fed females exhibited elevated insulin levels after refeeding following a 20 hr fast (**Supplementary Fig. 7C**). Furthermore, B6F on CR also exhibited a reduction in hepatic glucose production from alanine (**Supplementary Fig. 7D-G**). These results suggest that both male and female mice on a CR regimen adapt their metabolism to tightly maintain blood glucose levels, enabling them to face extended periods of daily fasting.

### The response of CR-fed mice to insulin is independent of time of feeding

Our initial observations suggested a 12-hr offset in insulin levels between AL- and CR-fed mice (**Fig. 2D-E**). This led us to hypothesize that plasma insulin levels might be influenced by the time of feeding, a theory supported by previous studies showing the circadian regulation of insulin secretion and its dependency on feeding schedule ^17,18^. To explore this further, we adjusted the feeding time for B6M to the onset of the dark cycle (ZT12) instead of the light cycle (ZT 1) and we examined the metabolic response to a fast (**Fig. 3A**). AL-fed control mice remained in their natural feeding state. For simplicity, we refer to our previous studies with B6M mice as Morning-Fed (CR mice fed ZT 0), and these new experiments as Night-Fed (CR mice fed diet at ZT 12).

**Figure 3.**
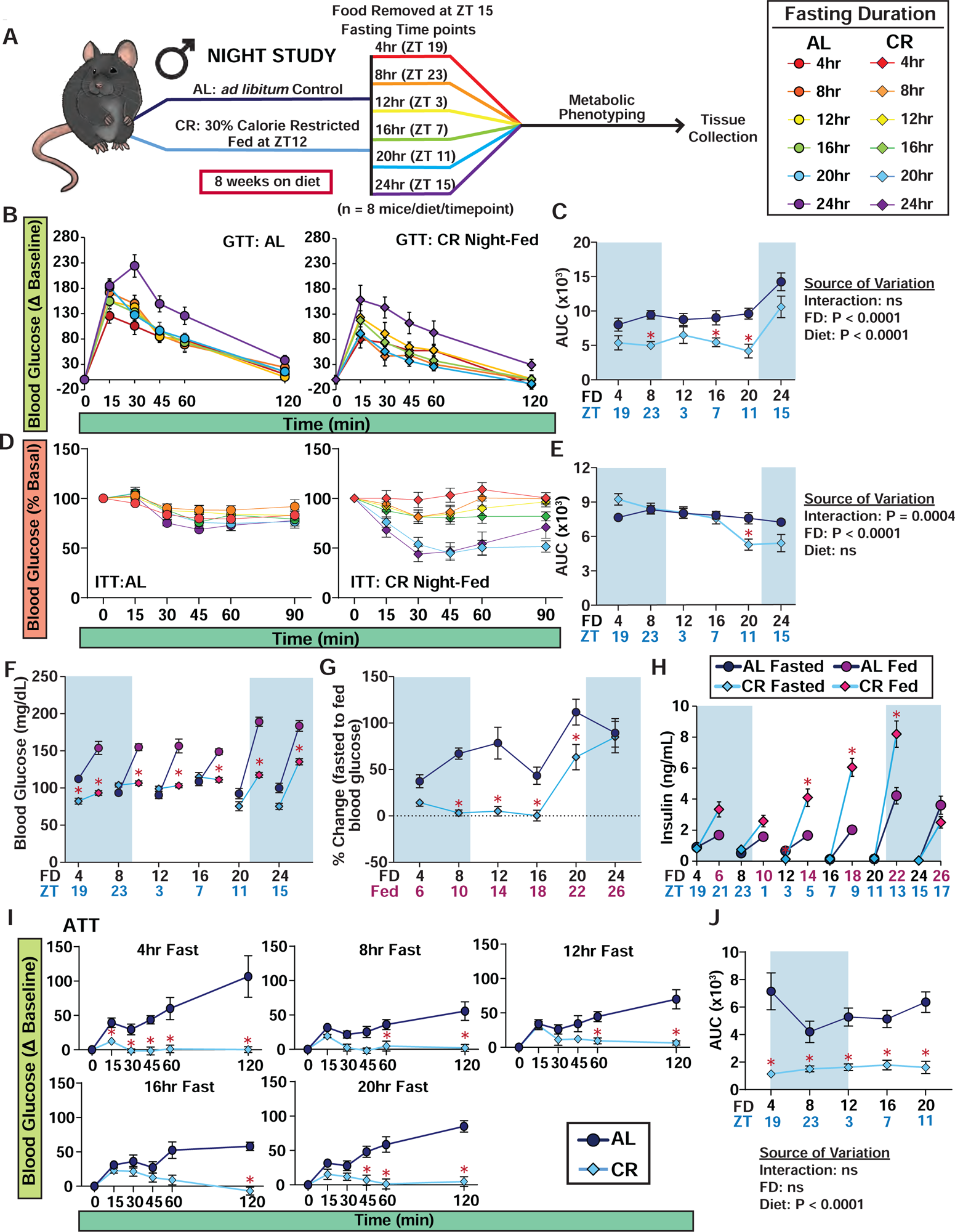
The response to insulin in CR feeding regimen is highly dependent on fasting duration and not time of feeding. (**A**) Experimental design of Night-Fed study. (**B-C**) Glucose tolerance tests of C57BL/6J male mice (**B**) and AUC (**C**); food was removed from all groups at ZT 15, and each group was then fasted for varying lengths of time. (**D-E**) Insulin tolerance tests of AL and CR-fed C57BL/6J male mice (**D**) and AUC (**E**). (**B-E**) 7-16 AL-fed and CR-fed biologically independent mice at each time point. (**C, E**) *p<0.05 AL-fed vs. CR-fed mice at each time point, Sidak’s test post two-way ANOVA. The overall effect of diet, fasting duration (FD), and the interaction represent the p-value from a two-way ANOVA. (**F-H**) Blood glucose level of mice fasted for 4, 8, 12, 16, 20 and 24hrs, prior to and subsequent to feeding (**F**), and percent change in blood glucose level from fasted to fed state (**G**). (**H**) Plasma insulin level of mice fasted for varying lengths of time, prior to and subsequent to feeding. (**F-H**) 7-15 AL-fed and CR-fed biologically independent mice at each time point. (**I-J**) Alanine tolerance tests of C57BL/6J male mice (**I**) and AUCs (**J**) (n=8 AL-fed and CR-fed biologically independent mice at each time point; the overall effect of diet, fasting duration (FD), and the interaction represent the p-value from a two-way ANOVA; *p<0.05 AL-fed vs. CR-fed mice at each time point, Sidak’s test post two-way ANOVA). Data represented as mean ± SEM. AUC, area under the curve. See **Supplementary Figures 7-9** for comparisons of AL and CR-fed mice at each time point.

In line with our Morning-Fed CR studies, the Night-Fed CR group also exhibited reduced weight, fat mass, and lean mass gain, with an overall effect of decreased adiposity (**Supplementary Figs. 8A-D**). Mice show a circadian pattern in their body weight, increasing during the active phase (i.e. dark cycle) and decreasing during inactive phase (i.e. light cycle)^19^, we evaluated body composition changes in AL- and CR-fed mice during a 24hr fast (**Supplementary Figs. 8E-H**). While there were no significant differences in body weight, fat mass and lean mass between the two groups during the fast, we observed a more rapid decrease in adiposity in CR-fed mice due to their lower absolute fat mass (**Supplementary Figs. 8E-H**).

Furthermore, like their Morning-Fed CR counterparts, Night-Fed CR mice had overall improved glucose tolerance compared to AL-fed mice, irrespective of the fasting duration, and this improvement reached statistical significance after 8, 16, or 20 hrs of fasting (**Figs. 3B-C**, **Supplementary Fig. 9**). Similar to what we observed with Morning-Fed CR mice, Night-Fed CR mice did not exhibit improved insulin sensitivity until fasting for at least 20 hours (**Figs. 3D-E, Supplementary Fig. 10**), suggesting that the response to insulin in CR-fed mice is dependent on fasting duration rather than the time of day.

We next measured fasted and fed blood glucose and serum insulin levels in the Night-Fed study (**Figs. 3F-H**). Initially, we found that Night-Fed CR mice maintained tight control over both their fasted and refed blood levels during the first 16 hours, similar to the Morning-Fed CR mice (**Figs. 3F-G**). However, contrary to our expectations of similar insulin levels between AL and CR-fed mice once feeding schedules were aligned, the Night-Fed CR mice showed a 2-3 fold increase in refed insulin levels after 12, 16, or 20 hours of fasting (**Fig. 3H, simplified graph in Supplementary Figs. 11A-B**). Lastly, examining hepatic glucose production post-alanine administration revealed that Night-Fed CR mice, like their Morning-Fed counterparts, suppressed glucose production from alanine (**Figs. 3I-J, Supplementary Fig. 11C-D**).

Collectively, these findings demonstrate that CR mice effectively resist changes in blood glucose induced by fasting and refeeding, irrespective of their feeding times. Importantly, while many of the metabolic effects of CR are linked to fasting duration, we find that insulin levels are primarily influenced by the feeding time rather than the fasting duration.

### Enhanced Blood Glucose Regulation in Night-Fed CR Mice During Refeeding

To assess potential differences between Morning-Fed and Night-fed CR mice, we calculated the Log_2_ ratio of CR to AL and then visualized these ratios using a radar chart (**Fig. 4A-D**). Our analysis revealed minimal variation in insulin and alanine responses between the Morning-Fed and Night-Fed groups (**Fig. 4B-C**). However, a notable shift in glucose tolerance was observed (**Fig. 4A**), as well as a significant difference seen in the percent change of blood glucose levels following refeeding.

**Figure 4.**
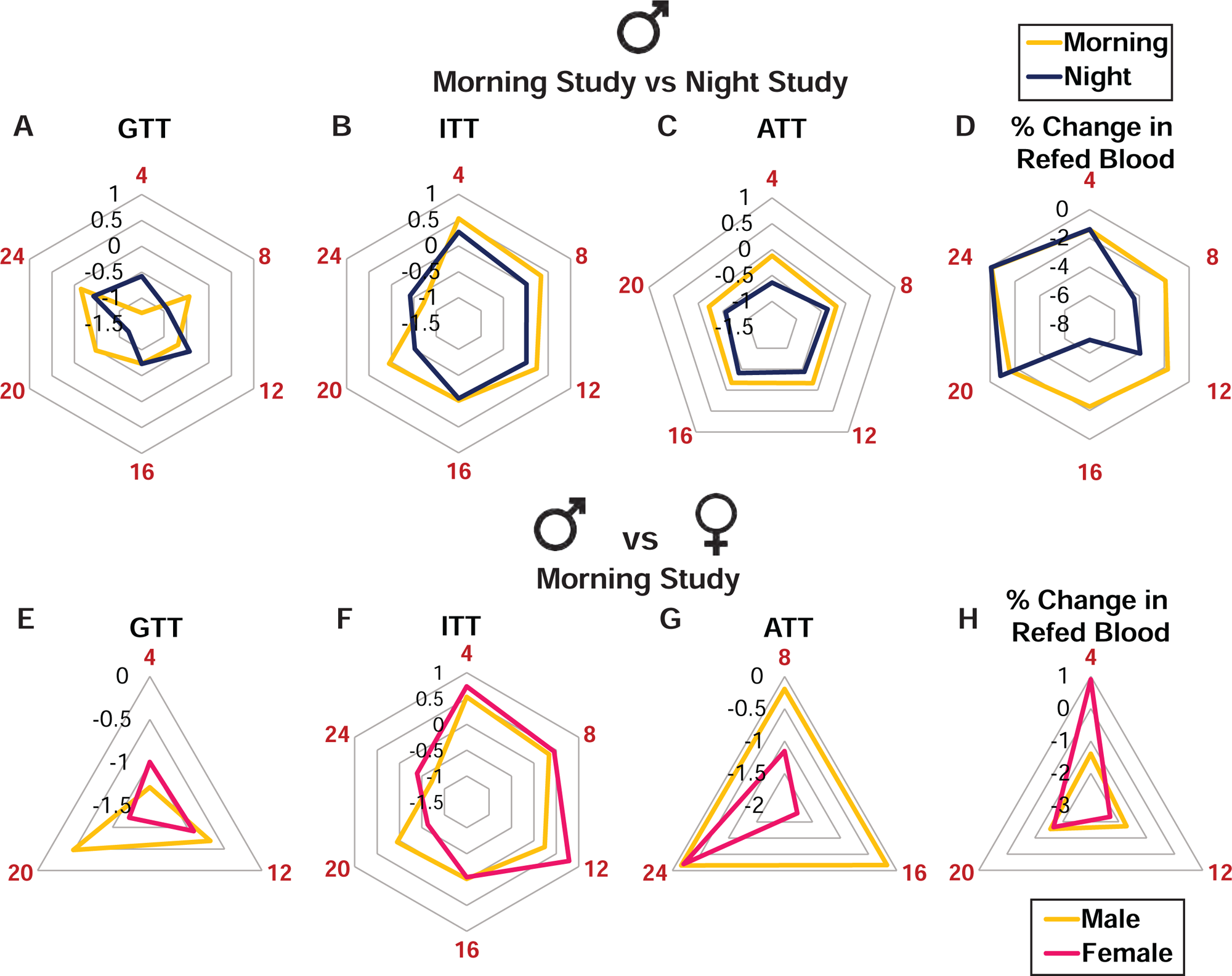
Comparison between the morning and night studies, and between males and females, related to Figures 1-3. Log_2_ ratio of CR to AL of AUC values calculated in Figures 1-3 and plotted on a radar chart. (**A-D**) Comparison between Morning and Night Study (A) GTT (B) ITT (C) ATT (D) % change in blood glucose level after mice are refed from fasted state during the refeeding study. (**E-H**) Comparison between males and females from the Morning Study (A) GTT (B) ITT (C) ATT (D) % change in blood glucose level after mice are refed from fasted state during the refeeding study. Red numbers represent fasting duration.

We next investigated the sex-dependent effects of CR on blood glucose regulation, again calculating the Log_2_ ratio of CR to AL and then visualizing the differences in both male and female mice using a radar chart (**Fig. 4E-H**). We noted some differences in the effect of CR on glucose and alanine tolerance between males and females. Specifically, CR improved glucose tolerance to a greater extent in males than females, whereas females showed a reduced glucose production from alanine (**Fig. 4E and G**). There was minor difference in the effect of CR on the response to insulin between male and female mice (**Fig. 4F**).

### AL and CR feeding have distinct and time-dependent effects on plasma metabolites and proteins

To gain additional insight into the effects of fasting and time-of-day on whole body metabolism, we used an untargeted metabolomics approach to examine plasma metabolite levels after either 8, 12, 16 or 24 hrs of fasting in AL- and CR-fed male mice from the Morning Study. We identified a total of 170 plasma metabolites; from this we utilized principal component analysis (PCA), plotting the first two principal components of each time point (**Fig. 5A**). Intriguingly, AL-fed and CR-fed mice displayed distinct metabolite profiles after 8 hours of fasting (blood collected at ZT 11) and 16 hours of fasting (blood collected at ZT 19). In contrast, the metabolite profiles of AL and CR mice were largely overlapping after either 12 hours or 24 hours of fasting (blood collected at ZT 15 and ZT3, respectively) (**Fig. 5A**).

**Figure 5.**
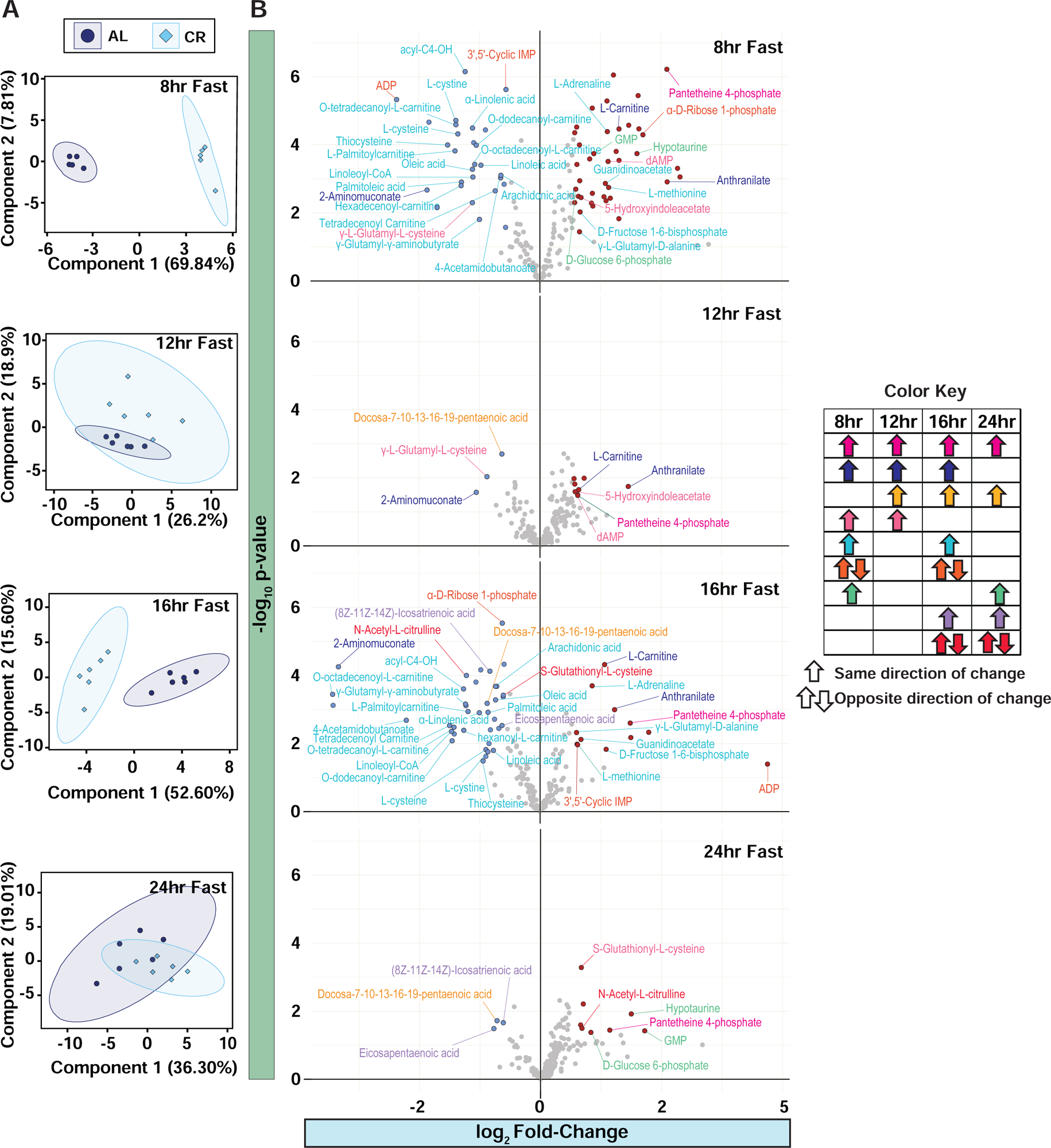
Plasma metabolite profile is dependent on time-of-day and diet. (**A**) Untargeted metabolomics were performed on the plasma of male C57BL/6J mice fed AL and CR diet from the Morning Study (n = 5-6 biologically independent mice per diet). PCA of plasma metabolites with AL and CR mice collected after an 8hr, 12hr, 16hr and 24hr fast. (**B**) Volcano plots of significantly increased plasma metabolites. Volcano plots show the statistical significance (p-value; y-axis) versus magnitude of change (fold-change (log_2_; x-axis). Significantly decreased metabolites are colored blue and significantly increased metabolites are colored red. Metabolite names are colored coded based on significant change between AL and CR at more than one time point (color key). See **Supplementary Figures 10-14** for comparisons of AL and CR-fed mice at each time point.

We next focused on metabolites that showed significant changes (p-value ≤ 0.05) at more than one time point (**Fig. 5B, Supplementary Table 1-2**). One standout finding was the consistent elevation of pantetheine-4-phosphate at all examined timepoints in CR-fed mice (**Fig. 5B**). Four metabolites were significantly altered at three of the fasting durations, with elevation of Anthranilate and L-Carnitine and decreased levels of 2-Aminomuconate; the remainder of the metabolites were altered only at two of the fasting durations (**Fig. 5B**).

While PCA and volcano plots offered a broad perspective of the significant changes associated with each diet and fasting durations, these analyses fall short in detecting trends that do not reach statistical significance. To bridge this gap, we analyzed significantly altered metabolites, aiming to discern the effects of diet, fasting duration and their interaction (**Supplementary Figs. 12-16**). We manually classified metabolites as involved in glucose metabolism (**Supplementary Fig. 12**), cysteine and methionine metabolism (**Supplementary Fig. 13**), arginine and tryptophan metabolism (**Supplementary Fig. 14**), mono- and poly-unsaturated fatty acid metabolism (**Supplementary Fig. 15**), and carnitine and nucleic acid metabolism (**Supplementary Fig. 16**).

Our analysis showed that in CR-fed mice, many of the glycolytic and TCA cycle metabolites were significantly increased, both overall and specifically at the 8hr time point (**Supplementary Fig. 12**). There were similar overall effects of a CR diet on the levels of the TCA cycle intermediates succinate, fumarate, and malate, as well as histidine, which is a α-ketoglutarate precursor. The pentose phosphate pathway metabolite α-D-Ribose-1-phosphate was significantly elevated after the 8, 16 and 24hr fasts, with a trend in elevation at the 12hr time point (**Supplementary Fig. 12**). In contrast to the glycolytic and TCA-cycle metabolites, we observed that CR-fed animals had an overall decrease in plasma levels of the branched chain amino acids (leucine, isoleucine, and valine), albeit with some interaction with fasting time (**Supplementary Fig. 12**).

Exploring further, we noticed trends in the cysteine sulfinic acid and transulfuration pathways in CR-fed mice, with most metabolites showing a decrease except for hypotaurine and γ-L-Glutamyl-D-Alanine (**Supplementary Fig. 13**). Contrasting patterns were observed in the levels of L-serine and L-methionine, which increased, while cysteine and cystine levels decreased in CR-fed mice (**Supplementary Fig. 13**). Interestingly, CR had a noticeable impact on tryptophan metabolism, including anthranilate, and significantly increased plasma levels of L-arginine and L-citrulline, metabolites in the urea cycle (**Supplementary Fig. 14**). Interestingly, CR-fed mice showed a general trend towards decreased levels of mono-unsaturated fatty acids (MUFAs) and poly-unsaturated fatty acids (PUFAs) (**Supplemental Fig. 15**). Conversely, medium and long-chain acyl-carnitine, short-chain acyl-carnitines and L-carnitine levels were elevated (**Supplemental Fig. 16**).

In parallel with the metabolomics study, we examined the plasma proteome of AL- and CR-fed animals, identifying a total of 294 proteins. Particularly at the 8 and 16 hr fasting period, distinct protein profiles were apparent between AL and CR-fed mice (**Supplementary Fig. 17 and Supplementary Tables 3-4**). However, to our surprise, less than 4% of proteins were significantly different between AL and CR-fed mice after 8 hours of fasting – with the differences becoming increasingly minimal as the fasting duration extended. Only a handful of proteins were significantly altered at more than one time point (**Supplementary Fig. 17**). Overall, our results suggest that CR feeding produces a unique rhythmic pattern in plasma metabolite levels without external nutrient influx, while circulating proteins are only minimally influenced by CR feeding.

### Hepatic and skeletal muscle mTORC1 activity is paradoxically not repressed by CR

The protein kinase mTORC1 (mechanistic Target Of Rapamycin) is a critical regulator of insulin sensitivity via S6K1-mediated feedback inhibition of insulin receptor substrate ^20,21^. Although it’s widely believed that CR promotes healthy aging by in part by reducing mTORC1 signaling^8^, this theory remains unproven. Indeed, CR and rapamycin induce non-overlapping and distinct molecular signatures in mice, suggesting that CR does not inhibit mTORC1 signaling. Furthermore, increased insulin sensitivity, generally reported as a characteristic of CR in mammals, might be predicted to paradoxically boost mTORC1 activity^22^. We hypothesized that high post-prandial mTORC1 signaling might explain the insulin resistance of CR-fed animals due to the negative feedback regulation of mTORC1 on insulin action.

To test our hypothesis, we sacrificed mice at multiple time points, matching the schedule from our *in vivo* metabolic phenotyping studies described above (**Figs. 6A-B**). We measured the phosphorylation of mTORC1 substrates and downstream readouts, as well as phosphorylation of the mTORC2 substrate AKT S473 and the PDK1 substrate AKT T308 in the liver (**Figs. 6C-D**) and skeletal muscle (**Figs. 6E-4F**). We found minimal differences in hepatic mTORC1 activity between AL- and CR-fed mice in either our Morning-Fed or Night-Fed studies (**Figs. 6C-D**). However, post-prandial S6 phosphorylation, a readout of mTORC1 activity, was higher in the liver of CR-fed mice than in AL-fed mice, reaching statistical significance in Morning-Fed CR mice at the 4 hour time point (**Figs. 6C-D**). In the skeletal muscle, we observed similar trends with heightened post-prandial S6 S240/S244 and S6K1 T389 phosphorylation in CR-fed mice at multiple time points (**Figs. 6E-F**). Additionally, the phosphorylation of other mTORC1 substrates, including 4E-BP1 T37/S46 and S757 ULK1, was elevated in the skeletal muscle of CR-fed mice at multiple time points (**Figs. 6E-F**).

**Figure 6.**
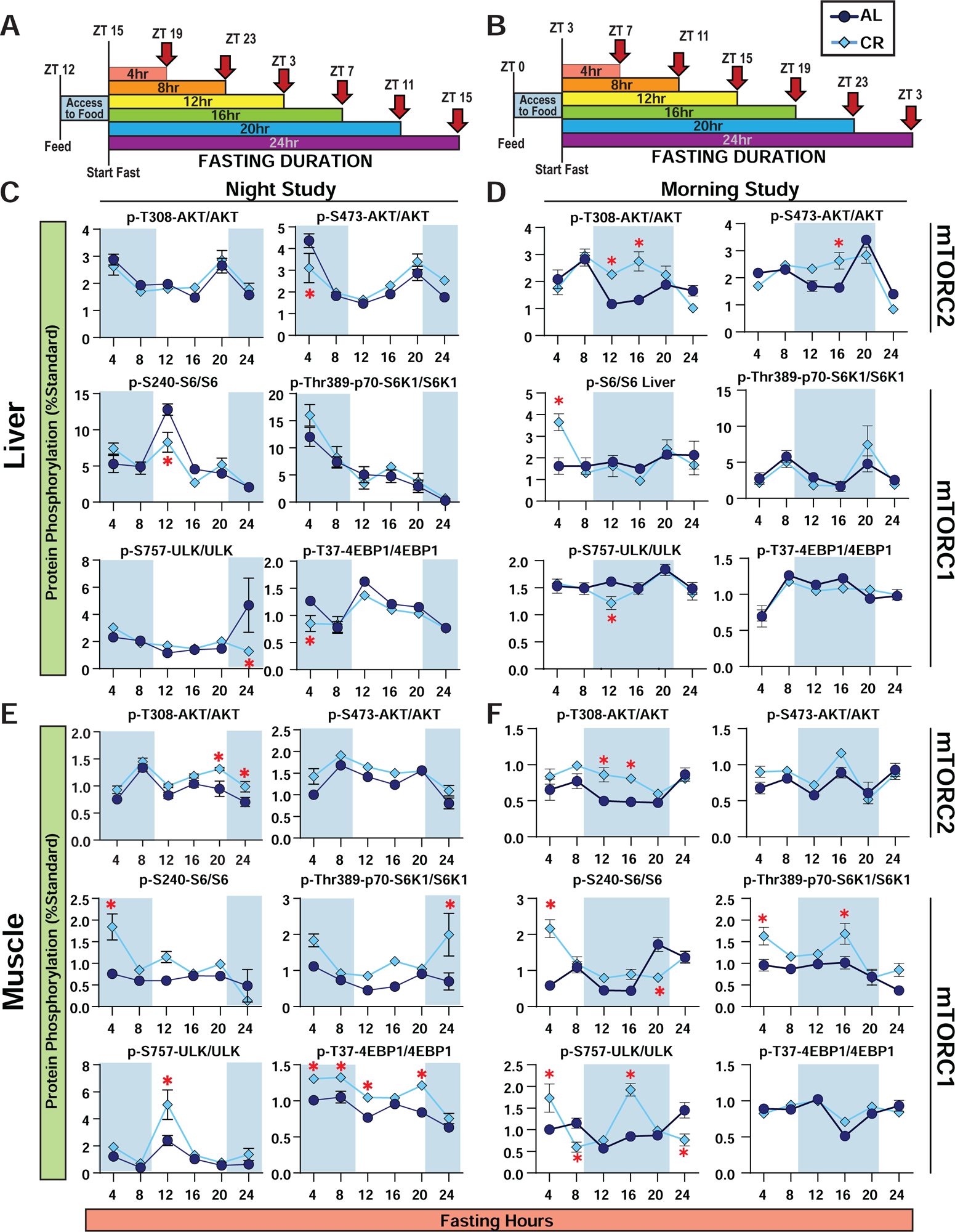
Liver and muscle mTORC1 activity is not constitutively repressed by CR. (**A-B**) Experimental design. Food was removed from Night-Fed mice at ZT15 (**A**) and Morning-Fed mice at ZT3 (**B**); groups of mice were sacrificed every 4 hours thereafter. (**C-D**) Liver protein lysate from Night-Fed (**C**) and Morning-Fed (**D**) mice was analyzed by Western blotting and quantified using ImageJ; y-axis reflects normalization of each phosphoresidue or protein to a liver standard run in duplicate on every gel (n=6 AL-fed and CR-fed biologically independent mice per time point; *p<0.05, Sidak’s test post 2-way ANOVA). (**E-F**) Muscle protein lysate from Night-Fed (**E**) and Morning-Fed (**F**) mice was analyzed by Western blotting and quantified using ImageJ; y-axis reflects normalization of each phosphoresidue or protein to a muscle standard run in duplicate on every gel (n=6 AL-fed and CR-fed biologically independent mice per time point; *p<0.05, Sidak’s test post 2-way ANOVA). Data represented as mean ± SEM.

The significant differences in phosphorylation of S6 in the liver and muscle at the earliest fasting timepoint (4hrs) (**Fig. 6C-F**) prompted us to consider that differences in mTOR signaling between AL- and CR-fed mice might be noticeable immediately after feeding. We synchronized the feeding times of all mice by fasting the mice at 6AM in the morning to encourage feeding, and then refeeding the mice starting at 6PM; we staggered the feeding allowing us to time the collection to the minute when each mice took its first bite (**Supplementary Fig. 18A**). When feeding was synchronized, AL and CR mice exhibited similar levels of mTORC1 activity in the liver (**Supplementary Fig. 18B**). Together, our results suggest that hepatic mTORC1 activity is not constitutively repressed in CR-fed mice, but instead results from the prolonged fasting between meals, and post-prandial mTORC1 activity is elevated in some tissues in CR-fed mice.

### Suppression of hepatic mTORC1 is required for CR-induced improvements to insulin administration in male mice

To investigate whether the suppression of hepatic mTORC1 activity is required to elicit the CR effect we utilized mice lacking hepatic *TSC1* (TSC1-LKO). TSC1-LKO mice are viable and have constitutively active hepatic mTORC1 signaling, even during fasting^23^, and we performed a series of metabolic measurements similar to those performed in Morning-Fed animals at two fasting durations (8hr and 21hr in males and females (**Figure 7**). There were no differences in glucose tolerance between the WT and TSC1-LKO mice within each respective diet for either males or females (**Figs. 7A and B**).

**Figure 7.**
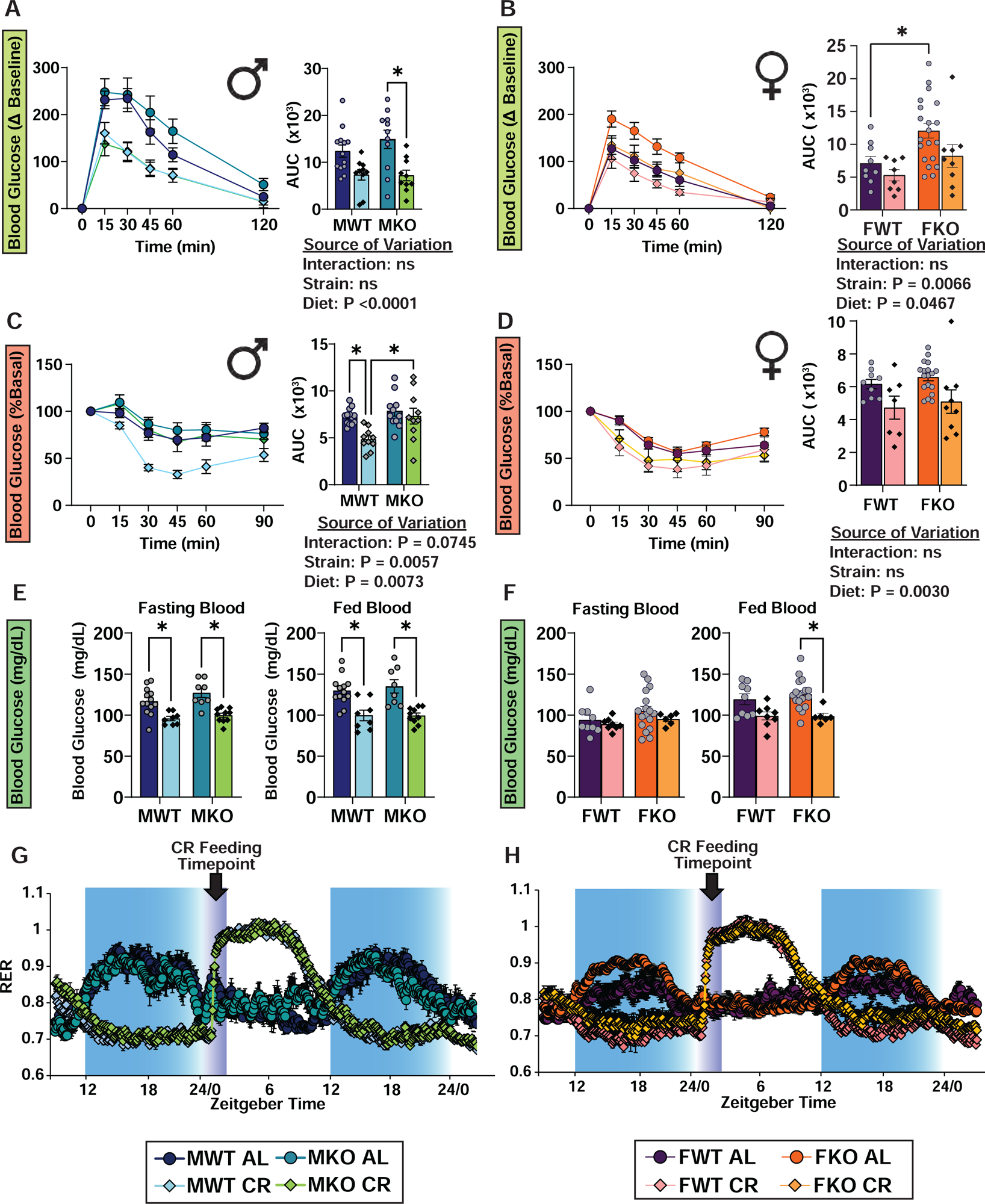
Constitutive suppression of hepatic mTORC1 activity is not required to produce the effect of CR. (**A-B**) Glucose tolerance tests in male (**A**) and female (**B**) mice lacking hepatic *Tsc1* (KO) and their wild-type (WT) littermates. (**C-D**) Insulin tolerance tests in male (**C**) and female (**D**) mice lacking hepatic *Tsc1* (KO) and their wild-type (WT) littermates. (**A,C**) MWT AL, n = 13; MWT CR, n = 10; MKO AL, n = 11; MKO CR, n = 10 biologically independent mice. (**B,D**) FWT AL, n = 9; FWT CR, n = 8; FKO AL, n = 20; FKO CR, n = 9 biologically independent mice. (**A-D**) statistics for the overall effects of genotype, diet, and the interaction represent the p value from a two-way ANOVA; *p<0.05, from a Sidak’s post-test examining the effect of parameters identified as significant in the 2-way ANOVA. (**E-F**) Blood glucose level of male (**E**) and female (**F**) mice fasted for 8hrs starting at ZT3 then refed at ZT12, and blood glucose level at ZT 14 after 2 hr refed state. (**E**) MWT AL, n = 13; MWT CR, n = 10; MKO AL, n = 11; MKO CR, n = 10 biologically independent mice. (**F**) FWT AL, n = 9; FWT CR, n = 8; FKO AL, n = 17; FKO CR, n = 6 biologically independent mice (**E-F**) *p<0.05 comparison between diet and genotype, Sidak’s test post two-way ANOVA. (**G-H**) Respiratory exchange ratio from metabolic chamber analysis of male (**G**) and female (**H**) mice. (**G**) MWT AL, n = 4; MWT CR, n = 4; MKO AL, n = 4; MKO CR, n = 4 biologically independent mice. (H) FWT AL, n = 4; FWT CR, n = 4; FKO AL, n = 4; FKO CR, n = 4 biologically independent mice. Data represented as mean ± SEM.

As expected, we found no significant differences in glucose tolerance or insulin sensitivity for either strain after an 8hr fast. In fact, response to insulin was significantly decreased by CR in TSC1-LKO mice of both sexes (**Supplementary Fig. 19A-D**). While CR diet significantly improved the response to insulin in WT males after a prolonged fast, unexpectedly we found that there was an interaction between diet and genotype (p=0.0745), with AL-fed and CR-fed TSC1-LKO male mice responding identically to insulin administration (**Fig. 7C**). In contrast, CR-fed female mice showed an improved response to insulin regardless of genotype (**Fig. 7D**). Furthermore, we found that both male and female mice fed a CR diet, regardless of genotype, maintained tight control of blood glucose levels even when challenged with refeeding (**Figs. 7E-F**). Lastly, we observed no effect of genotype on the CR-mediated changes in fuel utilization in either sex (**Figs. 7G-H**). Our data suggest that the metabolic effects of CR are largely independent of hepatic mTORC1 signaling. However, the effects of CR on whole body sensitivity to insulin administration are blocked by constitutive activation of hepatic mTORC1 in male mice.

### Aged mice have similar temporal responses to CR as young mice

Our previous studies showed that aged mice maintained on a CR diet from a young age retained a healthy phenotype^3^. Therefore, to examine if the variation in temporal conditions extended to aged animals, we examined aged male mice (22 months old) fed either an AL or CR diet starting at 4 months of age, fed in the morning (**Fig. 8A**). We found that as in young mice, aged CR-fed mice had improved glucose tolerance after 4-, 12-, and 20-hour fasts, but only showed improved response to insulin after a 20 hr fast (**Fig. 8B-C**). Furthermore, aged CR-fed mice similarly maintained tight control over both fasting and meal-stimulated blood glucose levels (**Fig. 8D-E**) and displayed a comparable trend in plasma levels of insulin as seen in younger mice **(Fig. 8F**). From these observations, we conclude that aged CR-fed mice respond similarly to young mice, underscoring the critical role of fasting duration in the apparent response to a CR diet.

**Figure 8.**
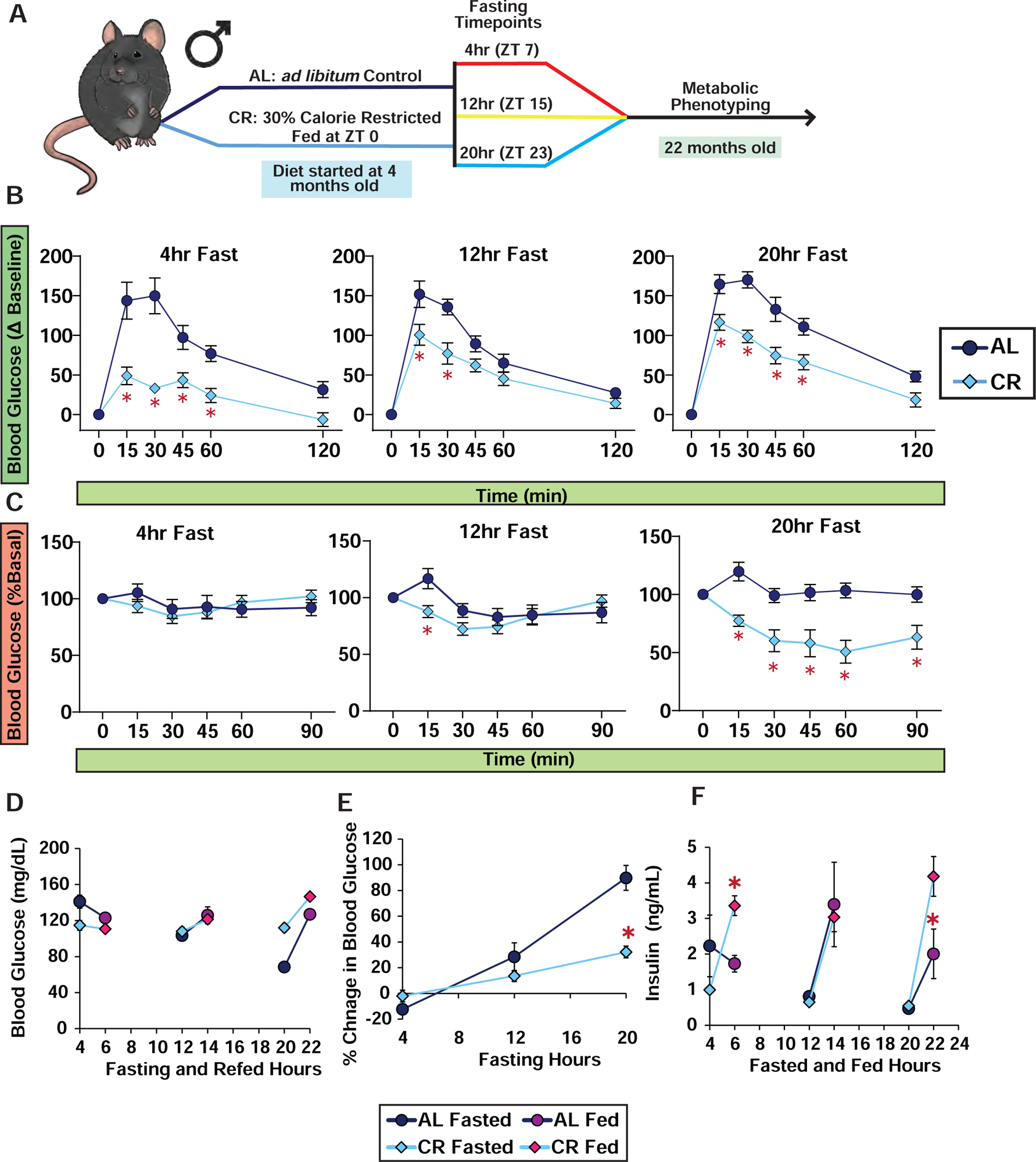
Morning-Fed aged male C57BL/6J mice have similar response to glucose and insulin as young male C57BL/6J. (**A**) Schematic of glucose and insulin tolerance tests for 22 month old mice. (**B**) Glucose tolerance tests of male C57BL/6J mice; food was removed from all groups at ZT3, and each group was then fasted for varying lengths of time. (**C**) Insulin tolerance test of male C57BL/6J mice; food was removed from all groups at ZT3, and each group was then fasted for varying lengths of time. (**D-E**) Blood glucose level of mice fasted for 4, 12, or 20 hrs, prior to and subsequent to feeding (**D**), and percent change in blood glucose level from fasted to fed state (**E**). (**F**) Plasma insulin level of mice fasted for varying lengths of time, prior to and subsequent to feeding. (**C-F**) n = 10 biologically independent mice per time point; *p<0.05 AL-fed vs. CR-fed mice at each time point, Sidak’s test post two-way ANOVA. Data represented as mean ± SEM.

## Discussion

There has been a long-standing interest in understanding the physiological and molecular basis by which CR functions to promote health and extend lifespan, with the eventual goal of finding small molecules or dietary regimens that can act as CR mimetics ^24,25^. Many signaling pathways have been proposed as key molecular mediators of the benefits of CR, but taken as a whole, our understanding of the mechanisms by which CR promotes lifespan and metabolic health remains the subject of debate. A widely used approach to understand how CR functions is to compare AL and CR-fed animals, identifying physiological and molecular differences between animals on each diet. This approach has shown that CR has a range of physiological effects, including improved insulin sensitivity ^26^ and protection from cancer ^27^ in many mammalian species, and modulates a host of signaling pathways. Surprisingly though, these beneficial effects on insulin sensitivity and cancer are each dispensable for the benefits of CR ^3,28,29^.

Why has this systematic approach to identifying mechanisms failed? CR feeding, at least in the context of once-per-day feeding, induces binge feeding behavior in which mice rapidly consume their daily allotment of food within 2-3 hours, resulting in a self-imposed daily fast until the next day’s feeding ^2,3,30^. A critical consequence of this collaterally imposed fast is that the feeding status of AL and CR-fed animals are not aligned, and thus, confounds our understanding of the physiological and molecular effects of CR. Furthermore, exposure to a daily fast could entrain an animal to develop an adaptive response to fasting which may confound the results of test preceded by fasting including GTTs and ITTs.

Here, we gained insight into how experimental variables – particularly, the length of the fasting period and the time of day of feeding – affect the observed metabolic response to CR. We show that the glucose tolerance of both AL and CR mice is dependent upon fasting duration, with glucose tolerance generally decreasing as we prolong the fast. Regardless of the length of the preceding fast, CR-fed mice had improved glucose tolerance relative to AL-fed mice in both morning and Night-Fed studies. However, we were surprised to discover that one of the key physiological hallmarks of CR in mammals – improved insulin sensitivity – was only observed in mice that had been fasted for at least 20 hours. In fact, CR-fed animals appeared to be insulin resistant when tested after a 4 hr fast, a response seen in both males and females and in both young and old mice. This response to insulin administration was not dependent upon time of day or time of feeding but was specifically dependent on how long the animals were fasted prior to the ITT. These results suggest that the widely conserved insulin-sensitizing effects of a CR diet are, at least to an extent, an artifact of how long mice have fasted prior to the assay. In particular, our results suggest that CR mice appear extraordinarily insulin sensitive relative to AL-fed controls due to the prolonged fasting time (16+ hours) routinely used, and suggest that under different paradigms, the literature would reflect that CR has no effect on the response to insulin. While our studies do not say that the insulin-sensitizing effects of CR are unimportant, they agree with our finding that CR can promote health and longevity in insulin-resistant mice ^29^.

Our assessment of glucose homeostasis was performed using GTTs and ITTs, and we recognize that there are alternative means to assess glycemic control and insulin sensitivity. In particular, we considered performing a hyperinsulinemic-euglycemic clamp, currently considered the gold standard to assess insulin sensitivity. However, this method is also subject to limitations, one of which is being able to observe the dynamic state that occurs under normal postprandial conditions ^31^. Furthermore, clamp data is biased towards detecting a muscle phenotype when performed in the prandial state ^32,33^. Nonetheless, the unique response to external glucose or insulin provides us with compelling evidence that AL and CR-fed mice respond differently to fasting.

To better understand the relationship between fasting duration and insulin sensitivity, we measured blood glucose and insulin levels following fasts of different length followed by a meal-stimulated insulin secretion (MSIS) test. While we expected a linear relationship between insulin levels and either fasting duration or diet regiments, we did not observe such trends. Rather, circulating insulin levels in CR-fed mice were higher than in AL-fed mice at some time points, and lower at others. While the interaction between time and insulin levels initially appeared random, we found that this was due to meal timing; shifting the fed insulin levels by 12 hours aligned AL and CR-morning fed mice, while feeding CR mice at night led to alignment of AL and CR-fed insulin values. Fasting insulin levels were dependent on fasting duration. Our results fit with previous research showing that insulin secretion is highly regulated by circadian controls and this mechanism is gated within a limited timeframe ^34,35^, and this response is altered depending on feeding schedule ^17,18^.

Our MSIS data gave us evidence for an adaptive response to fasting by CR mice, in which these mice were able to maintain tight control of blood glucose during a prolonged fast of up to 20 hrs, even after feeding. Our ITT data suggests that CR-fed mice were insulin resistant during this period. Therefore, we hypothesized that perhaps CR mice have adapted their metabolism to maintain tight blood glucose control independently of insulin. Addressing the role of insulin during a prolonged fast comes with many hurdles due to the difficulty of measuring the rate of the endogenous insulin secretion and the rate of insulin clearance. While at this time we could not test every aspect of the dynamics of insulin, we were able to inquire if insulin was required for the tight blood glucose control, we observed in CR-fed mice by acutely treating mice with diazoxide to suppress insulin secretion prior to a meal-stimulated insulin secretion test. To our surprise, blood glucose levels in CR mice rapidly rose to over 600 mg/dL, while AL mice showed only a moderate increase in blood glucose levels. These results revealed two key information: 1) insulin is indeed required for the maintenance of blood glucose level in CR-fed mice, acting to suppress glucose production via gluconeogenesis during a prolonged fast; and 2) CR feeding alters metabolism in order to preserve glucose during a fast. This data fits with our observation that CR-fed mice have reduced glucose production following alanine administration relative to AL-fed mice. Lastly, a speculation on our part, the decreased insulin sensitivity in CR-fed mice at the early fasting time points of the ITT, may be a negative feedback response to inhibit insulin sensitivity to maintain tight blood glucose levels during a prolonged fast.

To gain a broader perspective into what may be happening during a fast, we examined metabolite and protein levels during the Morning-Fed Study after an 8hr, 12hr, 16hr or 24hr fast. To our surprise, changes in metabolite levels did not have a linear relationship with fasting duration; vastly more metabolites were significantly altered after 8 and 16 hours of fasting than after 12 or 24 hours. Related to this, the majority of metabolites were significantly altered at only one or two time points. In addition to suggesting that there may be circadian or feeding-pattern dependent biological process affected by CR, our results highlight the possibility that a few key metabolites – such as pantetheine-4-phosphate – may serve as a timing independent serum marker of CR. Intriguingly, pantothenic acid (vitamin B5), a precursor of pantetheine-4-phosphate, has been found to be elevated in the blood of CR-fed mice in other studies ^36^. Together our data suggest that serum metabolome is greatly affected not only by CR, but also reflects the time of day and time since feeding when the samples were collected. Further work will be required to narrow down the specific contribution of diet, fasting duration, and alignment of feeding.

Recent work suggests that many of the observations we observed in our Morning Study may be the results of the different time discrepant feeding time of AL and CR mice. One study found that the timing of food intake plays an important role in the effect of CR on lifespan, with Night-Fed CR mice living longer than Morning-Fed CR mice ^37^. However, these results contrast with older studies showing no difference in lifespan between Morning-Fed and Night-Fed CR mice, as well as our own observation of a robust response to CR in Morning-Fed CR mice ^3,29^. We wondered if the time of day at which we fed mice would impact the response to fasting. Here, the only major difference we observed between Morning-Fed and Night-Fed CR mice was the insulin level at different times of the day. While we observed a similar trend in insulin level when feeding was aligned between AL- and CR-fed mice, there was a 2-4 fold increase in insulin level for Night-Fed CR mice. With the current studies we do not have a complete explanation for this observation. One possible explanation as to why we observe irregular insulin levels at different feeding times may be a dysregulated metabolism in Morning-Fed CR mice – which eat during the time mice would normally be sleeping, such as shift-workers who work and eat at night ^38,39^. More studies will be required to fully understand this phenomenon.

As we considered potential mechanisms for this unique response to insulin in CR-fed mice, an obvious choice was the mechanistic Target of Rapamycin (mTOR) pathway, which is a key regulator of lifespan and suppression of which has been proposed to be a key mediator of the effects of CR ^40–42^. mTOR is a protein kinase that is a key nutrient sensing hub, and importantly, a critical regulator of insulin sensitivity through S6K1 mediated feedback inhibition of insulin receptor substrate ^20,21,43^. While extensive work has been done using methods of genetic and pharmacogenetic inhibition of mTOR, research on how CR feeding regimen affects mTOR activity in mammals is scarce. Like others we initially hypothesized that mTORC1 activity was reduced in CR-fed animals, while AL-fed animals would have comparatively greater levels of mTORC1 activity. Surprisingly, this was not the case; hepatic mTORC1 activity was primarily mediated by fed and fasted state regardless of diet regimen, and this activity was perfectly matched when feeding was initiated around the same time (∼ZT 12). Furthermore, using mice with constitutively active mTORC1 (TSC1-LKO), we determined that suppression of hepatic mTORC1 was not required to elicit many of the phenotypes observed in CR animals. The nonlinear pattern of mTOR activity with fasting duration that we observed may be explained by previous works which have suggested that hepatic mTORC1 activity is regulated under circadian controls ^44–47^. Interestingly, we observed increased mTORC1 activity in the muscle, which may provide some evidence as to why CR mice eventually regain their lean mass. In addition, other studies have also found that CR, at least at specific time points and in specific cell types, acts to increase mTORC1 activity ^48,49^. Overall, these results suggest that inhibition of mTORC1, at least in the liver, may not be a key component of the CR response.

Our study was subject to limitations and may have contributed to what we observe in these studies. We acknowledge the limitations of GTT and ITT to assess glucose homeostasis as described above. The purpose of using these tests was to show the unique response to external stimulus based on fasting duration and timing of the day. Another limitation was the reliance of food consumption of the animals for our MSIS test, with CR-fed mice eating more food than the AL-fed mice (CR mice are capable of eating more food than AL animals at each timepoint). However, we did not see a trend in insulin levels based on the amount of food consumed; therefore, we do not think the food load contributed to the overall effect we observed here. Additionally, we did not measure insulin levels at earlier time points, which could have revealed effects of CR on the kinetics of insulin secretion. However, previous studies have shown that gastric emptying in mice takes about 74 minutes (with the completion of the meal) ^50^, and peak insulin levels after intraduodenal glucose delivery were observed after 120 minutes ^51^. Therefore, measuring insulin levels after 120 minutes should be sufficient to detect overall insulin secretion after a meal. Lastly, at this time we did not examine morning versus night fed in females or aged animals. More work will be required to understand the contribution of feeding time and pattern and fasting duration in the effects observed here in females and aged animals.

In conclusion, our work demonstrates that a number of the effects of a CR diet that have been studied for the last century are the result of (or, at a minimum, affected by) the specific temporal conditions of the assay chosen. As the best example, we discovered that CR-induced improvements in insulin sensitivity – an effect of CR which is believed to be conserved in all mammals – are only observed when an ITT is conducted following a prolonged fast. Indeed, we find that CR-fed mice demonstrate a strong post-prandial decrease in insulin sensitivity as assessed via IP administration of insulin. It will be interesting to learn if primates and humans on once-per-day CR regimens undergo similar daily periods of post-prandial insulin resistance and other metabolic shifts, and if the period of insulin sensitivity is limited to specific times of the day. One previous study with humans found a subset of participants on CR to have impaired glucose tolerance ^52^, and this may be due to differences in feeding pattern between the CR participants. We have previously shown that calorie restriction alone is not sufficient to produce the same response to insulin, and this once-per-day feeding was necessary in mice ^3^. And while other human studies have suggested the beneficial effects of fasting on health outcomes, controlled clinical trials remain to be explored ^53–55^.

Finally, while it is widely believed that CR promotes healthy aging in part by reducing activity of the protein kinase mTORC1, we have determined that when carefully aligning fasting and feeding periods, CR-fed mice do not show a global decrease in mTORC1 activity. Our results add to accumulating evidence that mTORC1 inhibition is not a major molecular mechanism by which CR promotes healthy aging and longevity. In yeast, deletion of *TOR1* is epistatic with CR; that is, CR does not further extend the lifespan of yeast lacking *TOR1* ^56^. Furthermore, the lifespan of flies is extended by the mTORC1 inhibitor rapamycin at every level of calorie intake ^57–62^, and extensive mice “omics” studies suggest that rapamycin and CR have distinct, largely non-overlapping effects ^22,63–66^, suggesting that CR and rapamycin do not function through the same molecular mechanism. In conclusion, our findings here demonstrate that the outcomes of CR experiments are heavily dependent upon the temporal conditions and how long it has been since both AL-fed and CR-fed mice have eaten, and we suggest that by taking these factors into account, researchers may now be able to truly understand how CR as well as other dietary interventions promote healthy aging.

## Materials and Methods

### Animals, Diets, and Feeding Regimens

All procedures were performed in conformance with institutional guidelines and were approved by the Institutional Animal Care and Use Committee of the William S. Middleton Memorial Veterans Hospital (Assurance ID: D16-00403) (Madison, WI, USA). Male and female C57BL/6J mice (stock number 000664) were purchased from The Jackson Laboratory (Bar Harbor, ME, USA) at 8 weeks of age and acclimated to the animal research facility for at least one week before entering studies. Liver specific *Tsc1* knockout mice (Tsc1-LKO) were generated by crossing mice expressing *Albumin-Cre* (stock number 003574) with mice expressing a conditional allele of *Tsc1* (*Tsc1^LoxP/LoxP^),* stock number 005680 as previously described ^23,67,68^. Aged male C57BL/6J mice were those used in our previously reported lifespan study ^3^. All animals were housed in a specific pathogen free (SPF) mouse facility with a 12:12 hour light/dark cycle maintained at 20°-22°C. All animals were placed on 2018 Teklad Global 18% Protein Rodent Diet for one week before randomization. Mice were randomized to either AL, *ad libitum* diet or CR, animals in which calories were restricted by 30% and fed once per day. Animals fed an AL and CR were fed 2018 Teklad Global 18% Protein Rodent Diet, Envigo Teklad. A stepwise reduction in food intake by increments of 10% per week, starting at 20% was carried out for mice in the CR group. Bodyweight and food intake were monitored weekly. Morning-Fed CR mice were fed daily at 6:00 - 7:00 a.m. and Night-Fed CR mice were fed daily at 6:00 – 7:00 p.m. The caloric intake of the mice in the AL group was calculated weekly to determine the appropriate number of calories to feed the mice in the CR group during the following week.

### Metabolic Phenotyping – Morning Fed

Glucose, insulin and alanine tolerance tests (GTT, ITT and ATT) were performed by feeding male mice at 6 a.m. and removing food at 9 a.m. We then tested the respective fasting durations: 4, 8, 12, 16, 20 or 24 hrs. At each time point, we injected a subset of mice with either glucose (1 g/kg), insulin (0.5U/kg) or alanine (2g/kg) intraperitoneally ^29^; no mouse was injected twice during the course of the 24 hr cycle. Glucose measurements were taken using a Bayer Contour blood glucose meter and test strips ^7^. Meal stimulated insulin tolerance (MSIS) test was performed by feeding mice at 6:00 a.m. and removing food at 9:00 a.m. We then measured and collected blood during their respective fasting durations: 4, 8, 12, 16, 20 or 24 hrs starting at 1 p.m. After the collection of fasted blood mice were allowed to feed for 2 hours which we then measured collected the fed blood. Insulin levels were assessed with Crystal Chem Insulin ELISA. Mouse body composition was determined using an EchoMRI Body Composition Analyzer. Adiposity was calculated by taking the fat mass and dividing by the sum of the fat mass and lean mass. For assay of multiple metabolic parameters (O_2_, CO_2_, food consumption and activity tracking), mice were acclimated to housing in a Columbus Instruments Oxymax/CLAMS metabolic chamber system overnight. AL-fed mice had access to food overnight and Morning-Fed CR mice were given their daily pellet the following day at 7 a.m. Then we removed food at 10 a.m. for both AL- and CR-fed mice and collected data from a continuous 24-hr period was then recorded and analyzed. Female mice were examined under similar conditions as male mice except ITTs were conducted on two separate weeks so that no mice were injected twice on a single day. Female mice were tested in the respective fasting durations – first week: 4, 12 or 20 hrs and second week: 8, 16, or 24hrs.

### Acute loss of insulin function with diazoxide

Acute diazoxide treatment study was performed by feeding mice at 6:00 a.m. and removing food at 9:00 a.m. We then injected diazoxide (0.75mg/bw) at 5:45 p.m. and then measured, collected fasted blood glucose, and allowed access to food for the mice. We initially set up the experiment for an MSIS (described above); however, the mice did not consume food. Therefore, we only measured blood glucose levels after 2 hr, 3hr and 4hr post-feeding.

### Metabolic Phenotyping – Night Fed

Glucose, insulin and alanine tolerance tests (GTT, ITT and ATT) were performed by feeding mice at 6:00 p.m. and removing food at 9:00 p.m. We then tested the respective fasting durations: 4, 8, 12, 16, 20 or 24 hrs and then injecting either glucose (1 g/kg), insulin (0.5U/kg) or alanine (2g/kg) intraperitoneally ^29^. Glucose measurements were taken using a Bayer Contour blood glucose meter and test strips. Meal stimulated insulin tolerance (MSIS) tests were performed by feeding mice at 6:00 p.m. and removing food at 9:00 p.m. We then measured and collected blood during their respective fasting durations: 4, 8, 12, 16, 20 or 24 hrs starting at 1 a.m. After the collection of fasted blood mice were allowed to feed for 2 hours which we then measured and collected the fed blood. Insulin levels were assessed with Crystal Chem Insulin ELISA. Mouse body composition was determined using an EchoMRI Body Composition Analyzer. For assay of multiple metabolic parameters (O_2_, CO_2_, food consumption and activity tracking), mice were acclimated to housing in a Columbus Instruments Oxymax/CLAMS metabolic chamber system for ∼24 hours. During this time AL-fed mice had continual access to food while CR mice were given their daily pellet at 6 p.m. Then we removed food at 9 p.m. for both AL- and CR-fed mice and collected data from a continuous 24 hr period was then recorded and analyzed.

### Metabolic Phenotyping – TSCKO

Glucose and insulin tolerance tests (GTT and ITT) were performed by feeding mice at 7 a.m. and removing food at 10 a.m. We then tested after either 8 or 21 hrs of fasting. At each time point, we injected a subset of mice with either glucose (1 g/kg or 2g/kg) or insulin (0.75U/kg for 8hr fast or 0.5U/kg for 21hr Fast) intraperitoneally ^29^.Glucose measurements were taken using a Bayer Contour blood glucose meter and test strips ^7^. Mouse body composition was determined using an EchoMRI Body Composition Analyzer. For assay of multiple metabolic parameters (O_2_, CO_2_, food consumption and activity tracking), mice were acclimated to housing in a Columbus Instruments Oxymax/CLAMS metabolic chamber system overnight. AL-fed mice had access to food overnight and Morning-Fed CR mice were given their daily pellet the following day at 7 a.m. and collected data from a continuous 24-hr period was then recorded and analyzed.

### Sacrifice and Collection of Tissues

Mice were sacrificed after 15 weeks on diet. Mice in the Morning-Fed studies were fed at 6 a.m. then food was removed starting at 9 a.m. and sacrificed in the respective fasting durations: 4, 8, 12, 16, 20 or 24 hrs. Mice in the Night-Fed studies were fed at 6 p.m. then food was removed starting at 9 p.m. and sacrificed in the respective fasting durations: 4, 8, 12, 16, 20 or 24 hrs. Following blood collection via submandibular bleeding, mice were euthanized by cervical dislocation and tissues (liver, muscle, iWAT, eWAT, BAT, and cecum) were rapidly collected, weighed, and then snap frozen in liquid nitrogen.

### Immunoblotting

Tissue samples from liver and muscle were lysed in cold RIPA buffer supplemented with phosphatase inhibitor and protease inhibitor cocktail tablets using a FastPrep 24 (M.P. Biomedicals) with bead-beating tubes (Stellar Scientific #BS-D1031-T20) and zirconium ceramic oxide bulk beads (VWR #10032-374). Protein lysates were then centrifuged at 13,300 rpm for 10 min and the supernatant was collected. Protein concentration was determined by Bradford (Pierce Biotechnology). 20 mg protein was separated by SDS-PAGE (sodium dodecyl sulfate-polyacrylamide gel electrophoresis) on 8%, 10%, or 16% resolving gels and transferred to PVDF membrane. To account for the variability between different western runs, we made a stock standard liver lysate sample which was run in duplicate on every gel. Membranes were blocked in 5% non-fat dry milk dissolved in TBST for 10 minutes and were then incubated in primary antibody diluted in 5% BSA (for phospho antibodies) or 5% milk (for non-phospho antibodies) overnight. The following commercial primary antibodies were used for immunoblot analysis p-S473 AKT (1:1,000; Cell Signaling Technology #4060), p-T308 AKT (1:1,000; Cell Signaling Technology # 2965), AKT (1:1,000; Cell Signaling Technology #4691), p-p70 S6K1 (1:500; Cell Signaling Technology #9234L), S6K (1:1,000; Cell Signaling Technology #2708S), p-S757-ULK (1:500; Cell Signaling Technology #5869S), ULK (1:500; Cell Signaling Technology #8054S), p-AMPK (1:1,000; Cell Signaling Technology #4188S), AMPK (1:1,000; Cell Signaling Technology #5831S), p-eIF2α (1:1,000; Cell Signaling Technology #3597L), eIF2α (1:1,000; Cell Signaling Technology #5324S), ATF4 (1:1,000; Cell Signaling Technology #11815S), HSP90 (1:1,000; Cell Signaling Technology #4877), p-S240/244 S6 ribosomal protein (1:1,000; Cell Signaling Technology #2215), S6 ribosomal protein (1:1,000; Cell Signaling Technology #2217), p-T37/S46 4E-BP1 (1:1,000; Cell Signaling Technology #2855), 4E-BP1 (1:1,000; Cell Signaling Technology #9452). Imaging was performed using a GE ImageQuant LAS 4000 imaging station (GE Healthcare). Quantification was performed by densitometry using NIH ImageJ software.

### Radar charts

Values for radar charters were calculated from the log_2_ fold-change from AL from each study and generated with Microsoft Excel. The distance from 0 represents the effect of CR versus AL-fed mice. Data are presented as the average of diet group per study (morning versus night or male versus females).

### Metabolomics

Plasma samples were thawed on ice then a 20 uL aliquot was treated with 480 uL of ice cold 5:3:2 MeOH:MeCN:water (v/v/v) then vortexed for 30 min at 4 degrees C. Supernatants were clarified by centrifugation (10 min, 12,000 g, 4 C). The resulting metabolite extracts were analyzed (20 uL per injection) by ultra-high-pressure liquid chromatography coupled to mass spectrometry (UHPLC-MS — Vanquish and Q Exactive, Thermo). Metabolites were resolved on a Kinetex C18 column (2.1 x 150 mm, 1.7 um) using a 5-minute gradient method in positive and negative ion modes (separate runs) exactly as previously described ^69^. Following data acquisition, .raw files were converted to .mzXML using RawConverter then metabolites assigned, and peaks integrated using Maven (Princeton University) in conjunction with the KEGG database and an in-house standard library. Quality control was assessed as using technical replicates run at beginning, end, and middle of each sequence as previously described ^70^.

### Proteomics

Plasma samples were digested in the S-Trap filter (Protifi, Huntington, NY). It was performed following the manufacturer’s procedure. Briefly, around 50 μg of plasma proteins were first mixed with 5% SDS. Samples were reduced with 10 mM DTT at 55 °C for 30 min, cooled to room temperature, and then alkylated with 25 mM iodoacetamide in the dark for 30 min. Afterward, to the samples was added a final concentration of 1.2% phosphoric acid and then six volumes of binding buffer (90% methanol; 100 mM triethylammonium bicarbonate, TEAB; pH 7.1). After gentle mixing, the protein solution was loaded to an S-Trap filter, spun at 2000 rpm for 1 min, and the flow-through collected and reloaded onto a filter. This step was repeated three times, and then the filter was washed with 200 μL of binding buffer 3 times. Finally, 1 μg of sequencing-grade trypsin and 150 μL of digestion buffer (50 mM TEAB) were added into the filter and digested at 47 °C for 1 h. The system was coupled to the timsTOF Pro mass spectrometer (Bruker Daltonics, Bremen, Germany) via the nano-electrospray ion source (Captive Spray, Bruker Daltonics) as previously describe ^71^. Raw data files conversion to peak lists in the MGF format, downstream identification, validation, filtering and quantification were managed using FragPipe version 13.0. MSFragger version 3.0 was used for database searches against a mouse database with decoys and common contaminants added. The identification settings were as follows: Trypsin, Specific, with a maximum of 2 missed cleavages, up to 2 isotope errors in precursor selection allowed for, 10.0 ppm as MS1 and 20.0 ppm as MS2 tolerances; fixed modifications: Carbamidomethylation of C (+57.021464 Da), variable modifications: Oxidation of M (+15.994915 Da), Acetylation of protein N-term (+42.010565 Da), Pyrolidone from peptide N-term Q or C (-17.026549 Da).

### Metabolomics and Proteomics Data Cleaning and Normalization

Plasma samples were analyzed via UHPLC-MS to determine plasma levels of 170 different metabolites, and plasma samples were analyzed via LC-MS/MS as described to determine levels of 856 distinct proteins. Metabolite peak intensity values that were equal to zero were removed from the dataset and missing data were imputed using a multiple imputation methodology (classification and regression tree (CART))^72^. Due to the large number of zero values in the proteomics dataset, the data were filtered so that data columns with >75% zeroes were dropped. Then the data were filtered further based on standard deviation (variables with SD <3 were dropped). After imputation of missing values and/or filtering, a log (base 10) transformation was applied to both the metabolomics and proteomics datasets.

Finally, the datasets were both scaled and centered using pareto scaling before further analyses were conducted. Final analysis of 170 metabolites and 294 proteins were conducted as described below. Fold-changes between each study group (CR-8hr/AL-8hr; CR-12hr/AL-12hr; CR-16hr/AL-16hr; CR-24hr/AL-24hr) were calculated using the raw data (non-normalized) for both the metabolomics and proteomics datasets. Then, Welch Two Sample t-tests were used to determine which metabolites/proteins were significantly different between each of the study groups. Volcano plots were generated to show significantly increased and decreased metabolites/proteins for each of the four study group pairs.

### Principal Components Analysis

Principal Components Analysis (PCA) was conducted to reduce dimensionality in the dataset for downstream analyses and to understand whether diet (AL vs. CR) and/or fasting duration (4, 8, 12, or 16 hours) resulted in distinct data clusters corresponding to specific metabolites/proteins. To determine which factors to retain, we used the standard Kaiser criterion, which suggests that factors with eigenvalues >1 should be extracted ^73^.

### Statistics

Data are presented as mean ± SEM unless otherwise specified. Analyses were performed using Excel (2010 and 2016, Microsoft) or Prism 9 (GraphPad Software). Statistical analyses were performed using one or two-way ANOVA followed by Sidak or Tukey-Kramer post hoc test specified in the figure legends. Other statistical details can also be found in the figure legends; in all figures, n represents the number of biologically independent animals. Sample sizes for metabolic studies were determined based on our previously published experimental results with the effects of dietary interventions ^74^, with the goal of having > 90% power to detect a change in area under the curve during a GTT (p<0.05). Data distribution was assumed to be normal, but this was not formally tested. All statistical analyses and data visualization of the metabolomics and proteomics data were performed using RStudio for Mac (RStudio Team [2022]. RStudio: Integrated Development for R. RStudio, Inc., Boston, MA URL). The alpha level for null hypothesis rejection was set at 0.05. Data are presented as mean ± standard error of the mean (SEM), unless otherwise noted.

### Randomization

All studies were performed on animals or tissues collected from animals. Animals of each sex and strain were randomized into groups of equivalent weight prior to the beginning of the *in vivo* studies.

### Data Availability

De-identified metabolomics and proteomics datasets are publicly available at the following dedicated GitHub repository: https://github.com/linsemanlab/Pak-et-al-2023. The complete datasets used for analyses and the source data used to generate each figure are included as part of this repository.

### Code Availability

The custom R code that was written for this publication is publicly available at the following dedicated GitHub repository: https://github.com/linsemanlab/Pak-et-al-2023.

## ACKNOWLEDGEMENTS

We would like to thank all members of the Lamming lab. The Lamming laboratory is supported in part by the NIH/NIA (AG056771, AG062328, AG081482, and AG084156 to D.W.L.), NIH/NIDDK (DK125859 to D.W.L.) and startup funds from the University of Wisconsin-Madison School of Medicine and Public Health and Department of Medicine to D.W.L. H.H.P. was supported by a NIA F31 predoctoral fellowship (AG066311). R.B. is supported by a training grant (T32DK007665). CLG was supported in part by Dalio Philanthropies, a Glenn Foundation for Medical Research Postdoctoral Fellowship, and by grant HF-AGE AGE-009 from Hevolution Foundation to CLG. Support for this research was provided by the University of Wisconsin - Madison Office of the Vice Chancellor for Research and Graduate Education with funding from the Wisconsin Alumni Research Foundation. This work was supported in part by the U.S. Department of Veterans Affairs (I01-BX004031 and IS1-BX005524), and this work was supported using facilities and resources from the William S. Middleton Memorial Veterans Hospital. The Paredes laboratory was supported in part by the Movement Disorder Foundation. The content is solely the responsibility of the authors and does not necessarily represent the official views of the NIH. This work does not represent the views of the Department of Veterans Affairs or the United States Government.

## AUTHOR CONTRIBUTIONS

All authors participated in the performance of the experiments and/or analyzed the data. HHP, ANG, DAP and DWL prepared the manuscript.

---

**Supplementary Figure 1.**
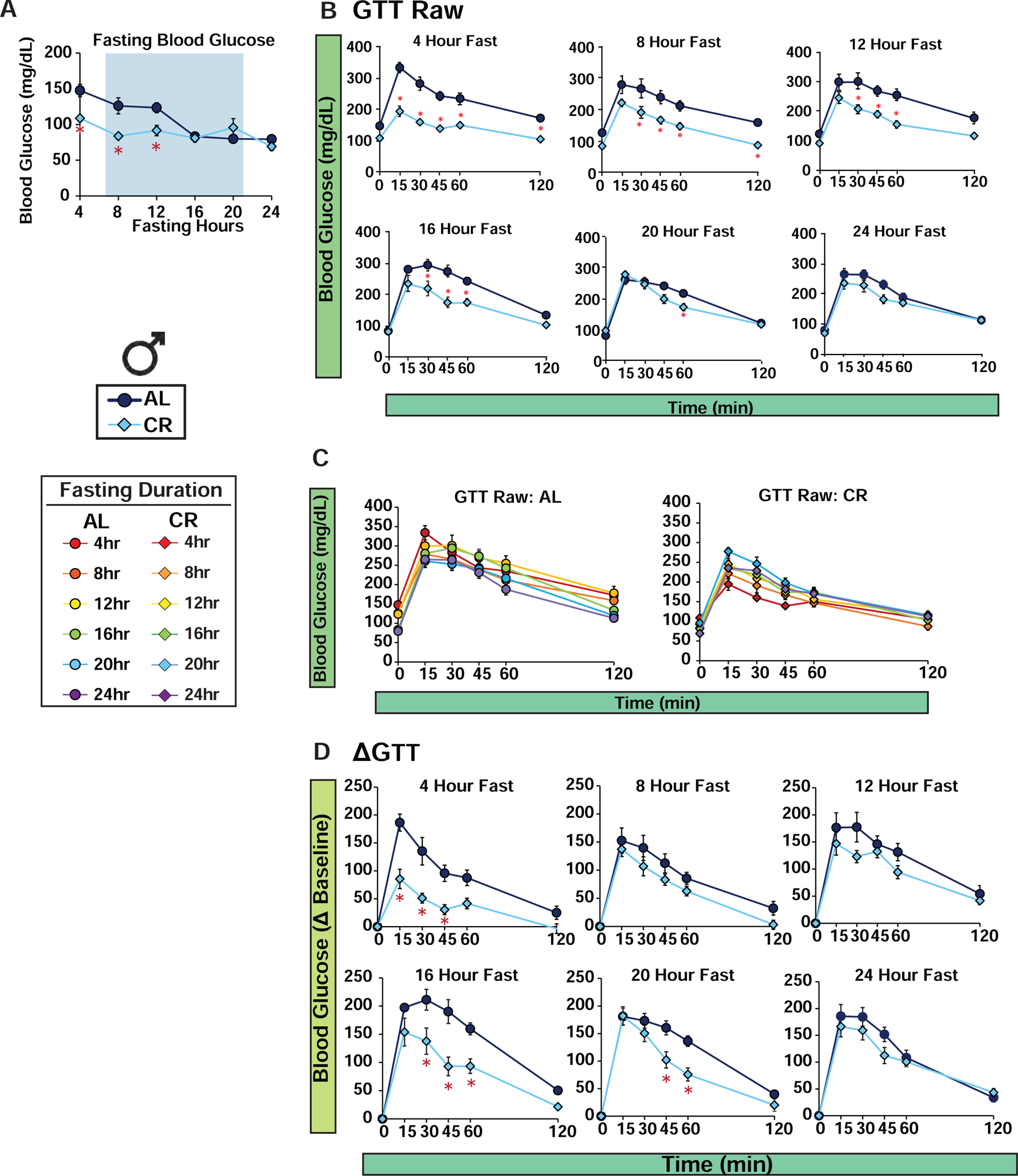
Raw and delta values for glucose tolerance test in Morning-Fed males, related to Figure 1. (**A**) Fasting blood glucose prior to GTT. (**B**) Raw GTT values. (**C**) Combined raw GTT for AL and CR. (**D**) Delta values for GTT. AL, n = 7-8 per time point; CR, n = 7-8 per time point biologically independent mice; *p<0.05 AL-fed vs. CR-fed mice at each time point, Sidak’s test post two-way repeated measures ANOVA. Data represented as mean ± SEM.

**Supplementary Figure 2.**
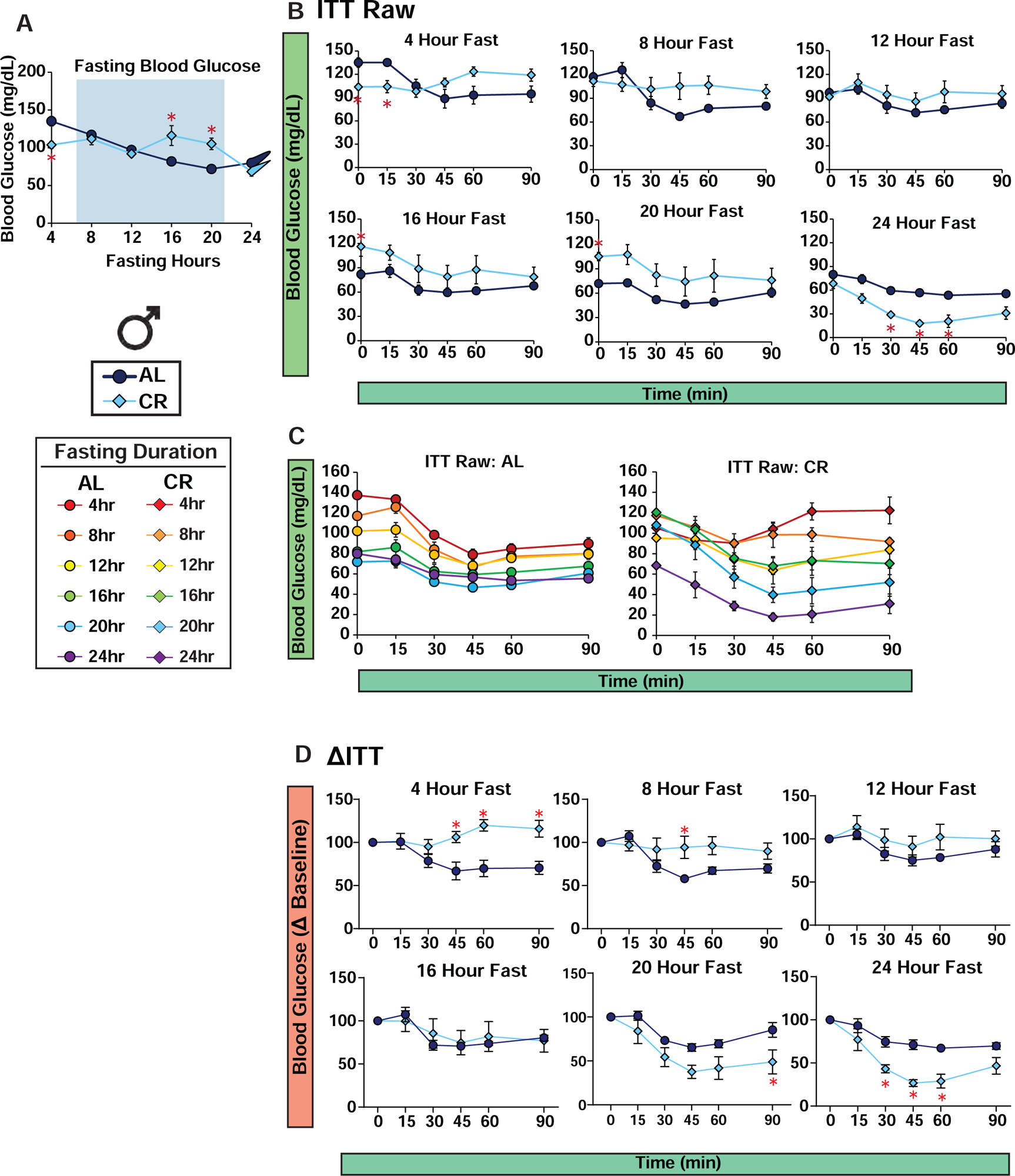
Raw and delta values for insulin tolerance test in Morning-Fed males, related to Figure 1. (**A**) Fasting blood glucose prior to ITT. (**B**) Raw values for ITT (**C**) Combined raw ITT for AL and CR. (**D**) Delta values for ITT (AL, n = 7-8 per time point; CR, n = 7-8 per time point biologically independent mice; *p<0.05 AL-fed vs. CR-fed mice at each time point, Sidak’s test post two-way repeated measures ANOVA. Data represented as mean ± SEM.

**Supplementary Figure 3.**
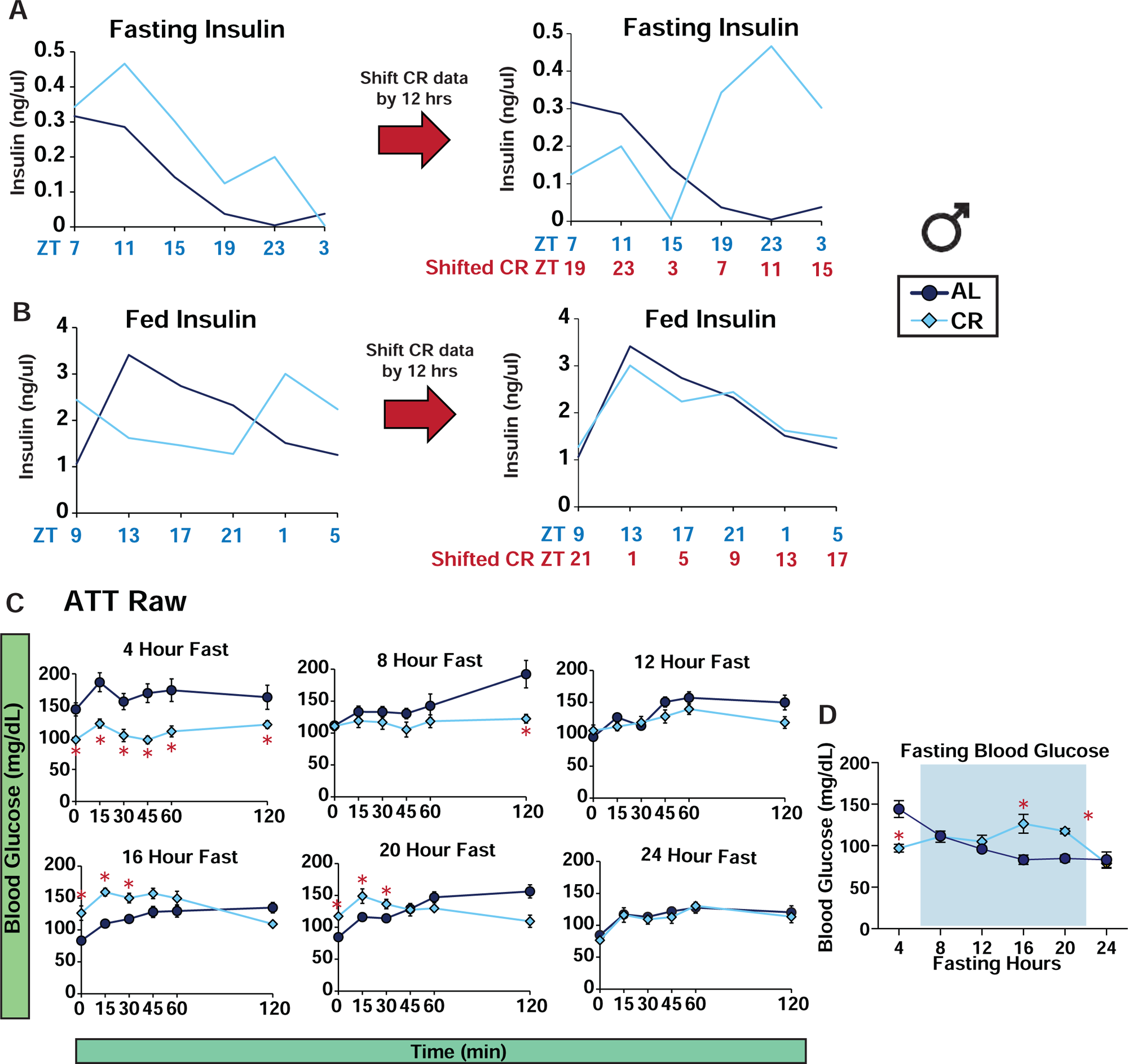
Raw values for alanine tolerance test in Morning-Fed males, related to Figure 2. (**A**) Simplified representation of fasted insulin level related to Fig. 2D-E. (**B**) Simplified representation of fed insulin level related to Fig. 2D-E. (**C**) Raw values for ATT with (**D**) fasting blood glucose. (AL, n = 7-8 per time point; CR, n = 7-8 per time point biologically independent mice; *p<0.05 AL-fed vs. CR-fed mice at each time point, Sidak’s test post two-way repeated measures ANOVA. Data represented as mean ± SEM.

**Supplementary Figure 4.**
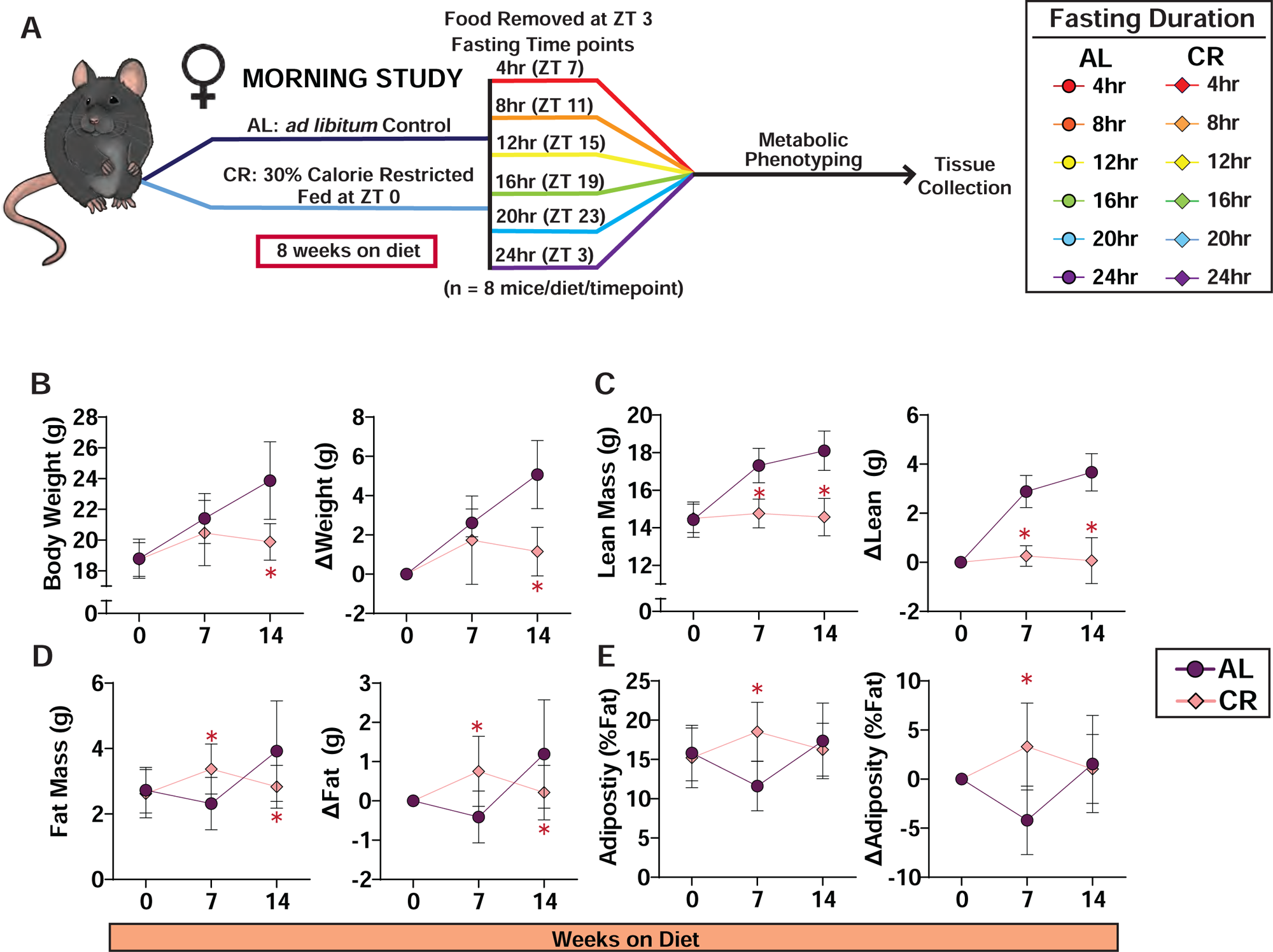
Body composition of female C57BL/6J mice in Morning-Fed study. (**A**) Experimental design of female C57BL/6J mice under Morning-Fed study. (**B-E**) Body composition measurement of female C57BL/6J mice under Morning-Fed conditions (AL, n = 7-8 and CR, n = 7-8 biologically independent mice) *p<0.05 AL-fed vs. CR-fed mice at each time point, Sidak’s test post two-way repeated measures ANOVA. Data represented as mean ± SEM. (**A**) Total body weight with change in body weight from baseline. (**B**) Fat mass with change in fat mass from baseline. (**C**) Lean mass with change in fat mass from baseline. (**D**) Adiposity with change in adiposity from baseline.

**Supplementary Figure 5.**
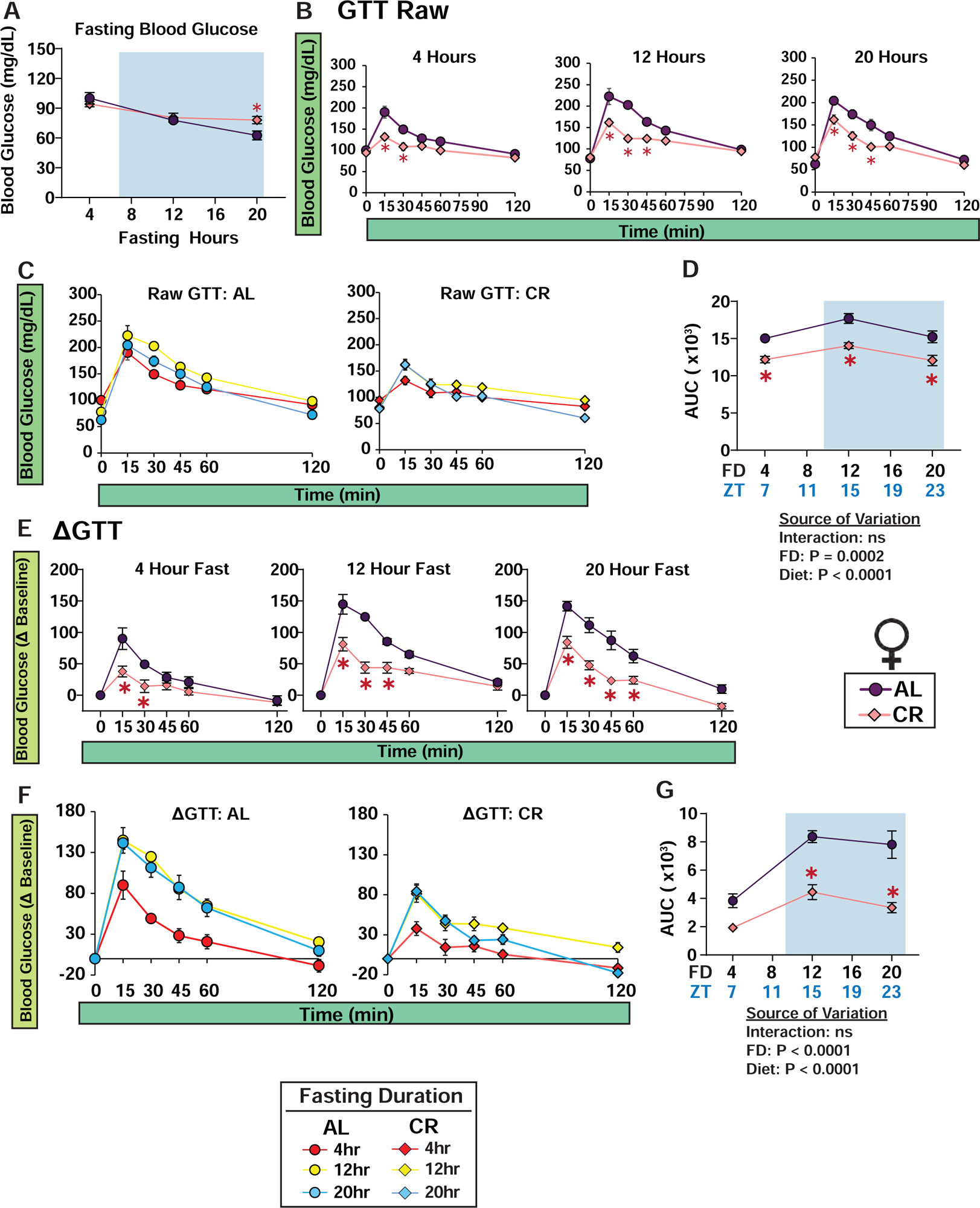
Glucose tolerance test of female C57BL/6J mice in Morning-Fed Study. (**A**) Fasting blood glucose prior to GTT. (**B**) Raw GTT values. (**C**) Combined raw GTT for AL and CR (**D**) with AUC. (**E**) Delta values for GTT. (**F**) Combined delta GTT for AL and CR (**G**) with AUCAL, n = 7-8 per time point; CR, n = 7-8 per time point biologically independent mice; *p<0.05 AL-fed vs. CR-fed mice at each time point, Sidak’s test post two-way repeated measures ANOVA. Data represented as mean ± SEM.

**Supplementary Figure 6.**
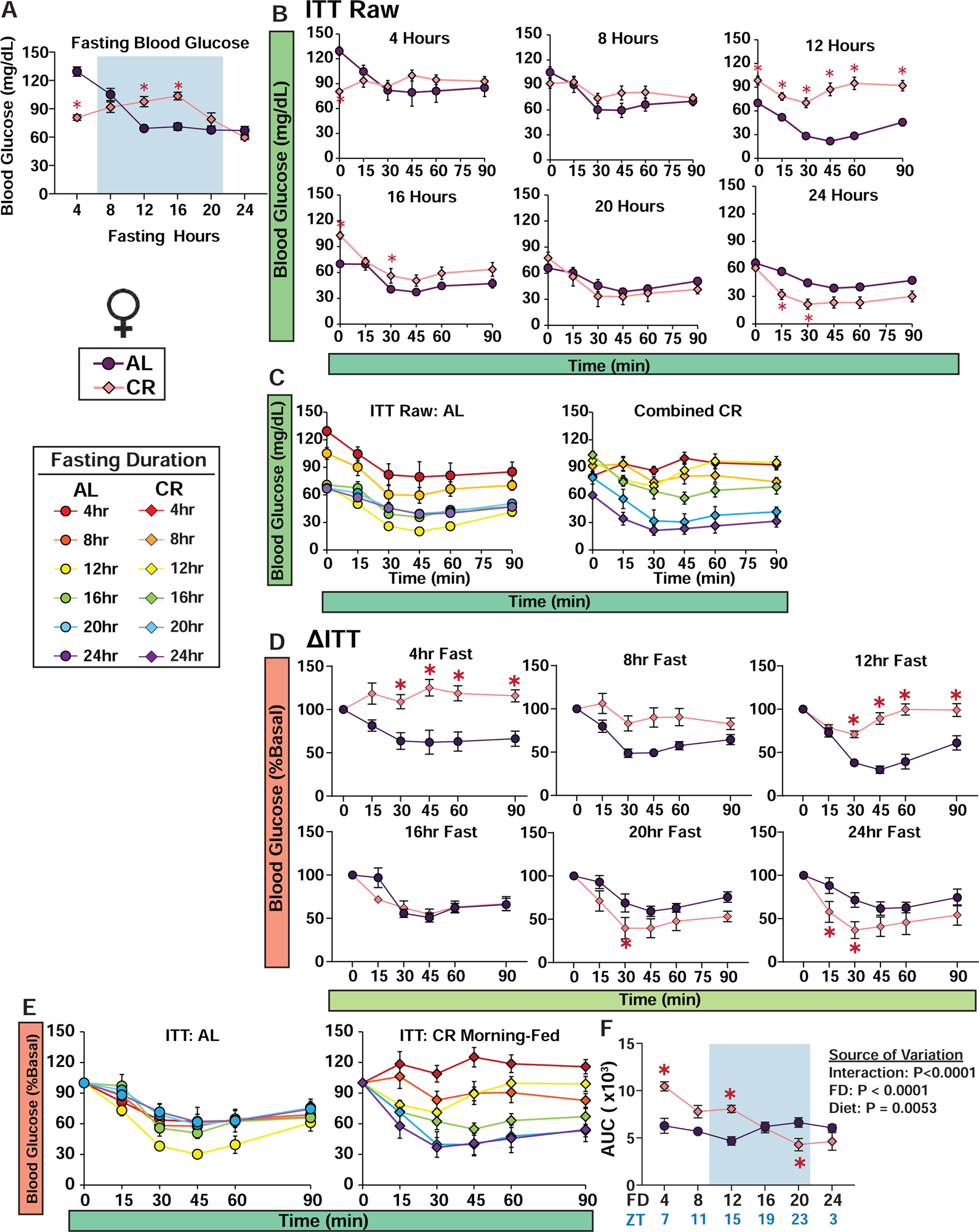
Insulin tolerance test of female C57BL/6J mice in Morning-Fed Study. (**A**) Fasting blood glucose prior to ITT. (**B**) Raw ITT values. (**C**) Combined raw ITT for AL and CR. (**D**) Delta values for GTT. (**E**) Combined delta GTT for AL and CR (**F**) with AUCAL, n = 7-8 per time point; CR, n = 7-8 per time point biologically independent mice; *p<0.05 AL-fed vs. CR-fed mice at each time point, Sidak’s test post two-way repeated measures ANOVA. Data represented as mean ± SEM.

**Supplementary Figure 7.**
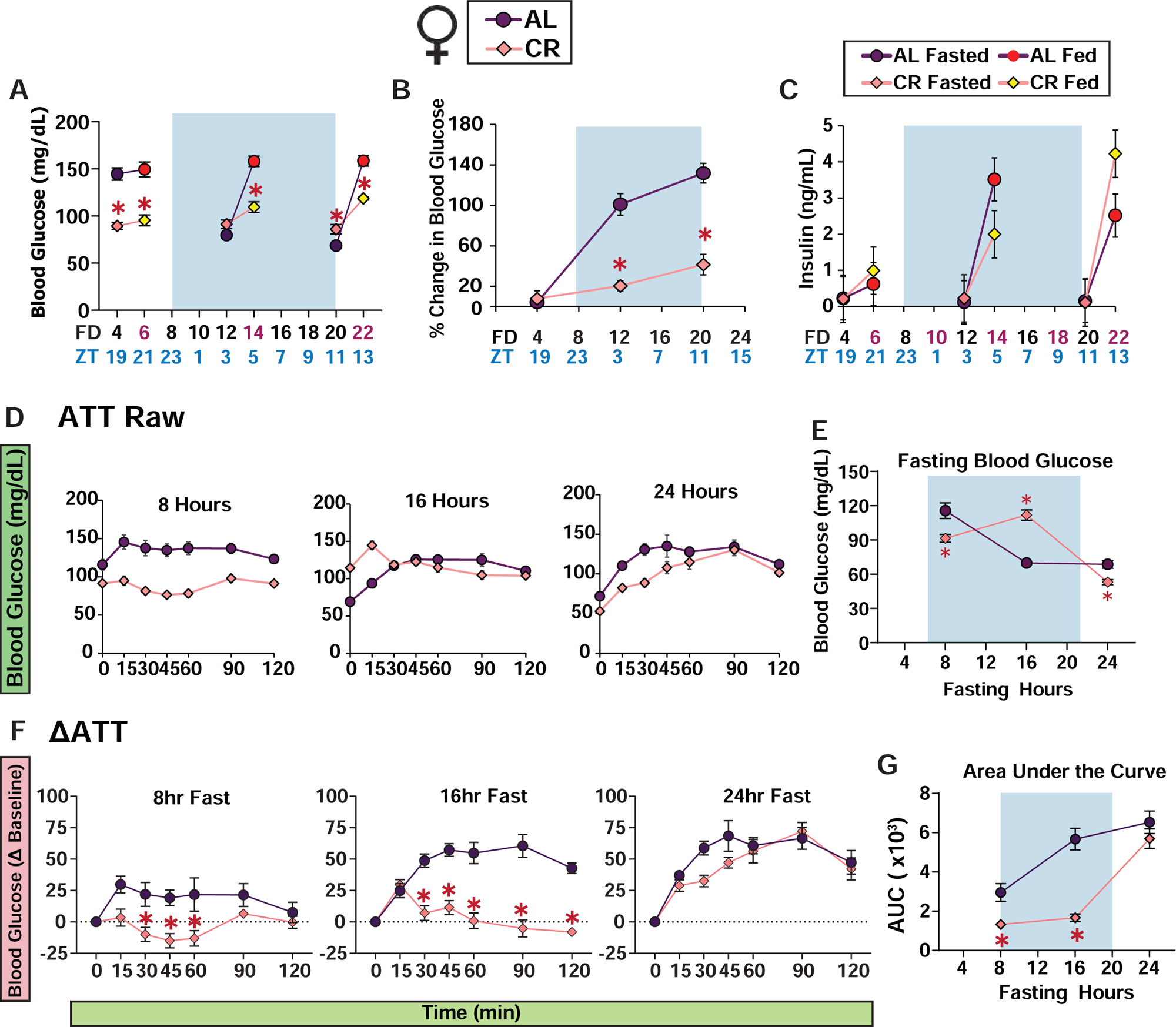
Meal-stimulated insulin level of female C57BL/6J mice in Morning-Fed study. (**A-B**) Blood glucose level of fasted and fed state during respective fasting timepoints (**A**), percent change in blood glucose level from fasted to fed state (**B**). (**C**) Insulin level of fasted and fed state mice during respective fasting/refed timepoints. (**D**) Raw values for ATT with (**E**) fasting blood glucose. (**F**) Delta values for ATT and (**G**) AUC (**A-G**) AL, n = 7-16 per timepoint; CR, n = 7-16 per timepoint, biologically independent mice; *p<0.05 AL-fed vs. CR-fed mice at each time point, Sidak’s test post two-way repeated measures ANOVA. Data represented as mean ± SEM.

**Supplementary Figure 8.**
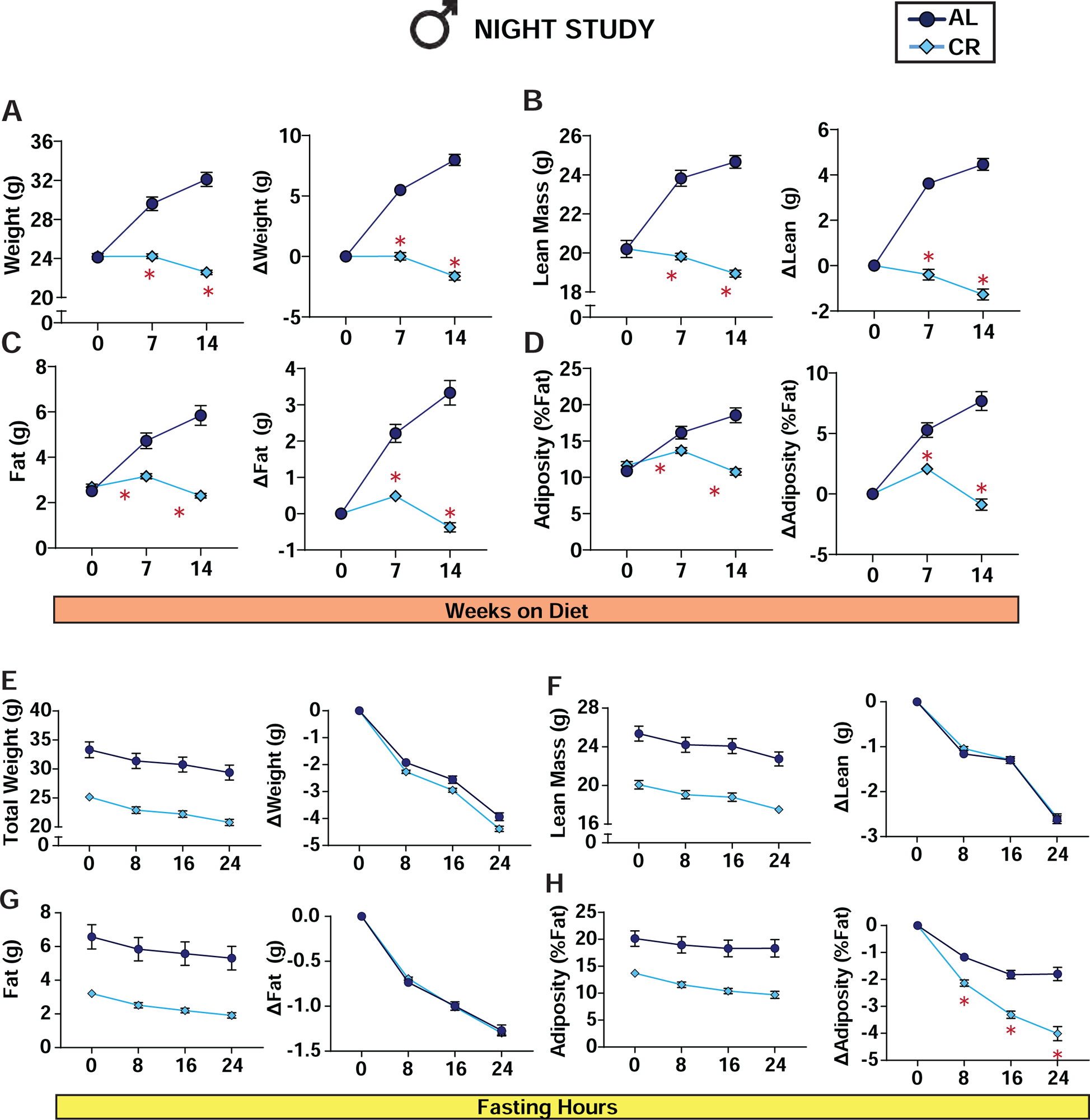
Body composition of Night Study male C57BL/6J mice and change in body composition during a 24hr fast, related to Figure 3. (**A**) Total body weight with % change in body weight from baseline. (**B**) Lean mass with % change in lean mass from baseline. (**C**) Fat mass with % change in fat mass from baseline. (**D**) adiposity with % change in adiposity from baseline. (**A-D**) AL, n = 27 and CR, n = 27 biologically independent mice. *p<0.05 AL-fed vs. CR-fed mice at each time point, Sidak’s test post two-way repeated measures ANOVA. Data represented as mean ± SEM. **(E-H)** Mice were fed at ZT 12, and body composition was measured starting at ZT 15 (0hr fasted). All mice were fasted after the initial baseline measurements and body composition was measured again at ZT 20 (8hr fasted), ZT 4 (16hr fasted), and ZT 12 (24hr fasted). (**E**) Total body weight with % change in body weight from baseline, (**F**) lean mass with % change in lean mass from baseline (**G**), fat mass with % change in fat mass from baseline, (**H**) adiposity with % change in adiposity from baseline. (**F-H**) AL, n = 12 and CR, n = 12 biologically independent mice. *p<0.05 AL-fed vs. CR-fed mice at each time point, Sidak’s test post two-way repeated measures ANOVA. Data represented as mean ± SEM.

**Supplementary Figure 9.**
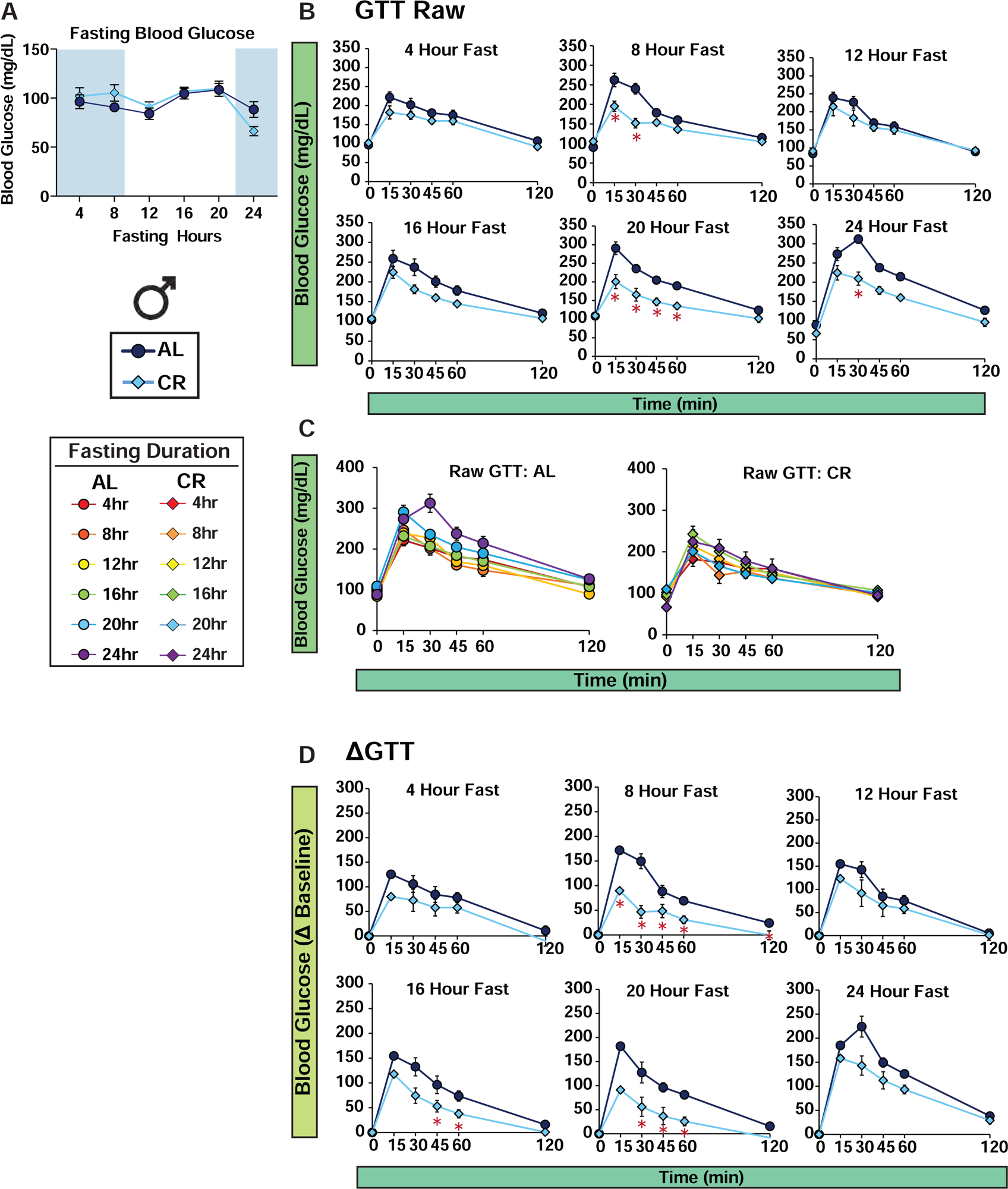
Raw and delta values for glucose tolerance test of C57BL/6J mice in Night-Fed study, related to Figure 3. (**A**) Fasting blood glucose prior to GTT. (**B**) Raw GTT values. (**C**) Combined raw GTT for AL and CR (**D**) Delta values for GTT. (AL, n = 7-16 per time point; CR, n = 7-16 per time point biologically independent mice; *p<0.05 AL-fed vs. CR-fed mice at each time point, Sidak’s test post two-way repeated measures ANOVA. Data represented as mean ± SEM.

**Supplementary Figure 10.**
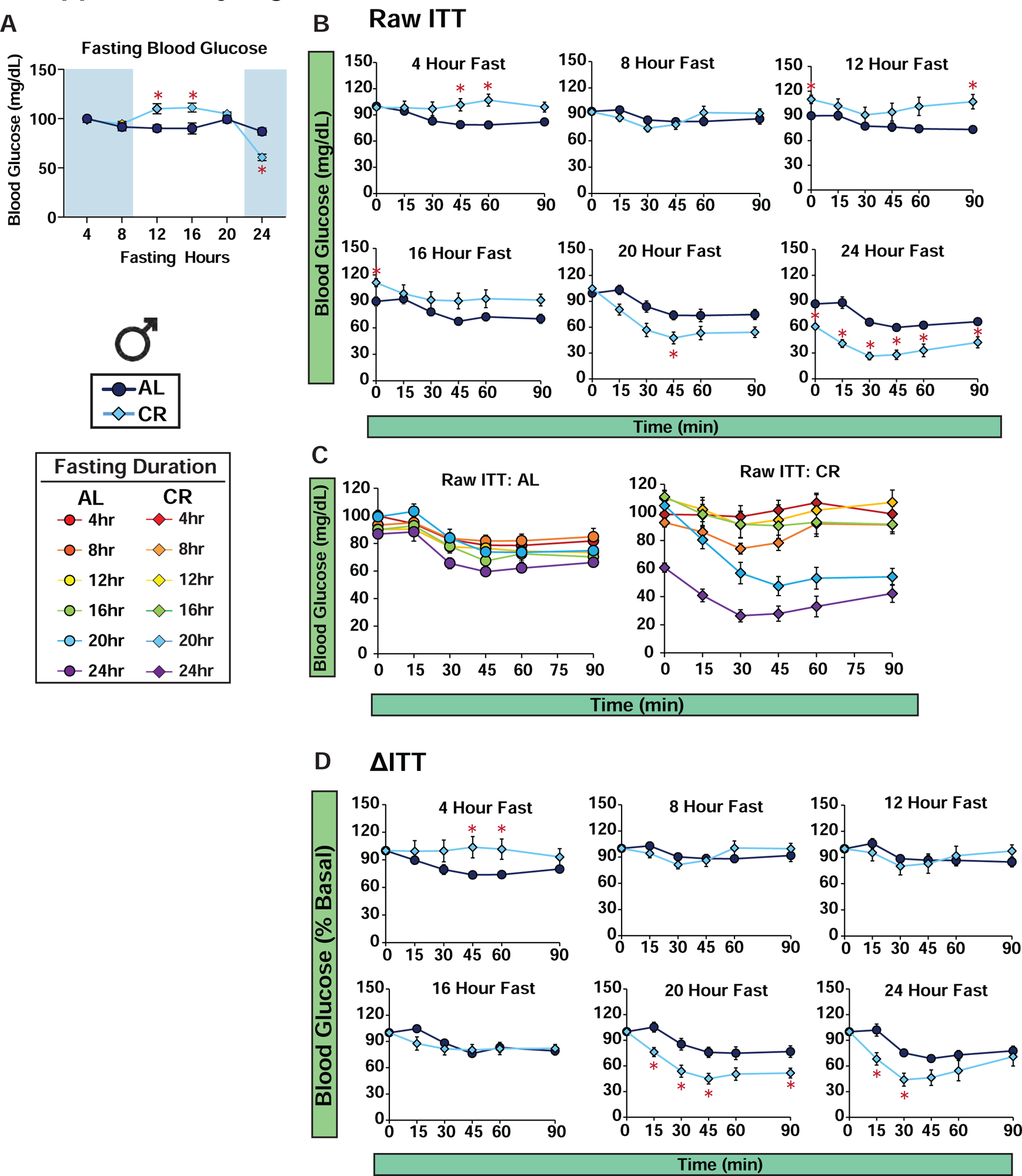
Raw and delta values for insulin tolerance test of C57BL/6J mice in Night-Fed study, related to Figure 3. (**A**) Fasting blood glucose prior to ITT. (**B**) Raw ITT values. (**C**) Combined raw ITT for AL and CR (**D**) Delta values for ITT. (AL, n = 7-16 per time point; CR, n = 7-16 per time point biologically independent mice; *p<0.05 AL-fed vs. CR-fed mice at each time point, Sidak’s test post two-way repeated measures ANOVA. Data represented as mean ± SEM.

**Supplementary Figure 11.**
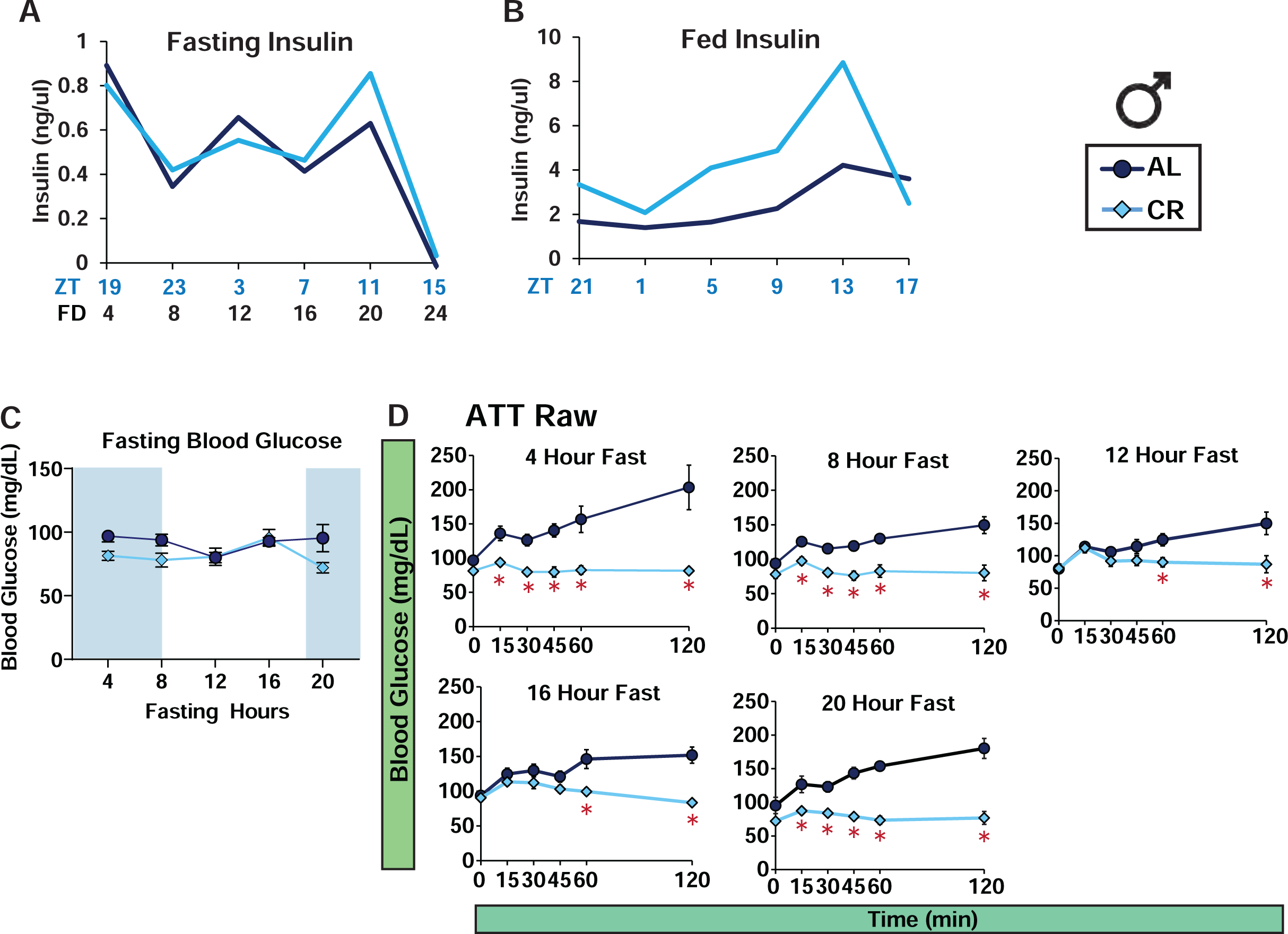
Raw values for alanine tolerance test in Night-Fed males, related to Figure 3. (**A**) Simplified representation of fasted insulin level related to Fig. 3H. (**B**) Simplified representation of fed insulin level related to Fig. 3H. (**C**) Fasting blood glucose prior to ATT. (**D**) Raw values for ATT. (AL, n = 7-16 per time point; CR, n = 7-8 per time point biologically independent mice; *p<0.05 AL-fed vs. CR-fed mice at each time point, Sidak’s test post two-way repeated measures ANOVA. Data represented as mean ± SEM.

**Supplementary Figure 12.**
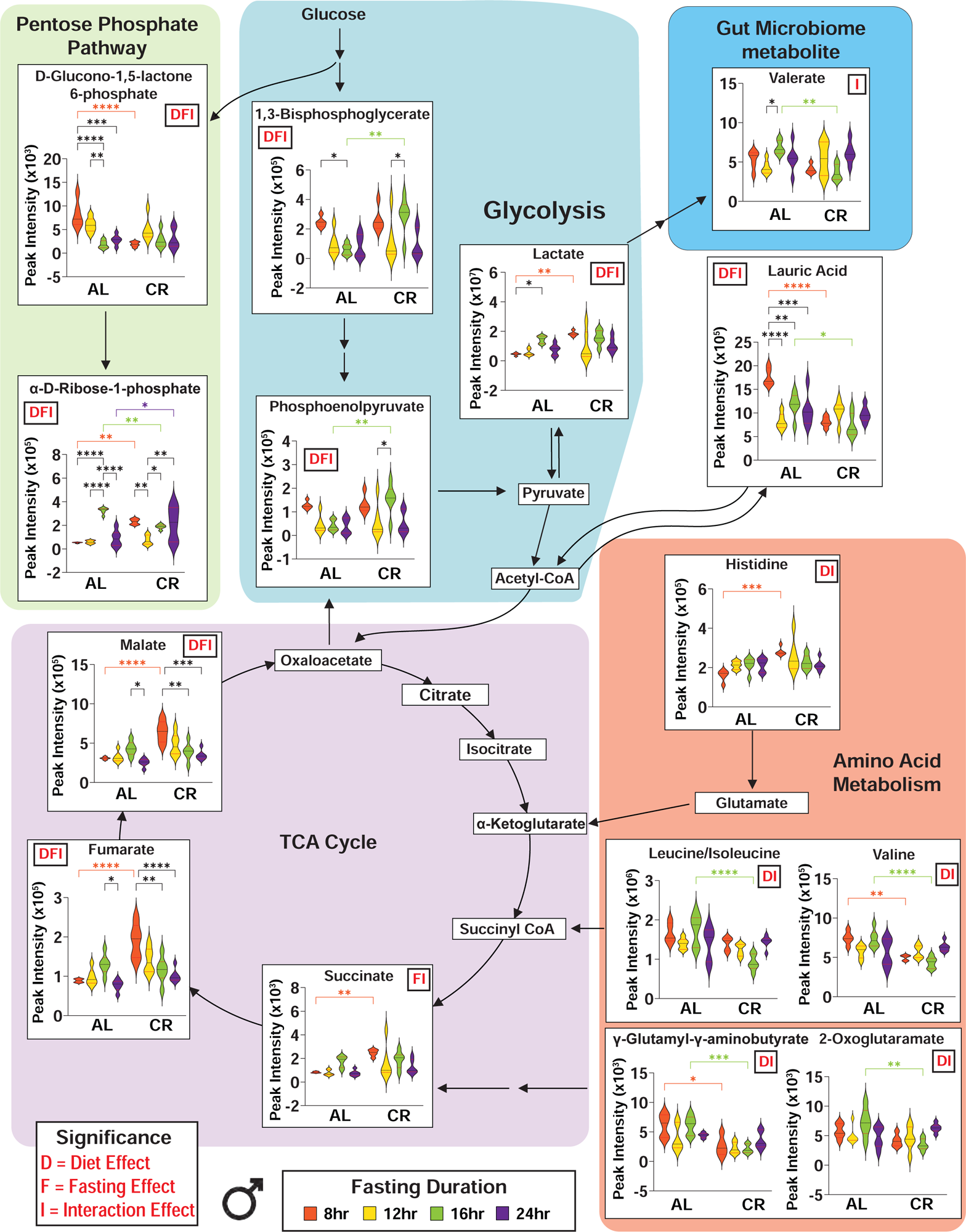
Plasma metabolite data involved in pentose phosphate pathway, glycolysis, TCA cycle and Amino Acid from the untargeted metabolomics analysis, related to Figure 4. *p<0.05 Tukey’s test post two-way ANOVA; D/F/I indicate a significant main effect of diet (D), fasting duration (F), or the diet and fasting duration interaction from the two-way ANOVA(I). The overlaid violin plot shows center as median (black lines) and 25^th^-75^th^ percentiles (red lines).

**Supplementary Figure 13.**
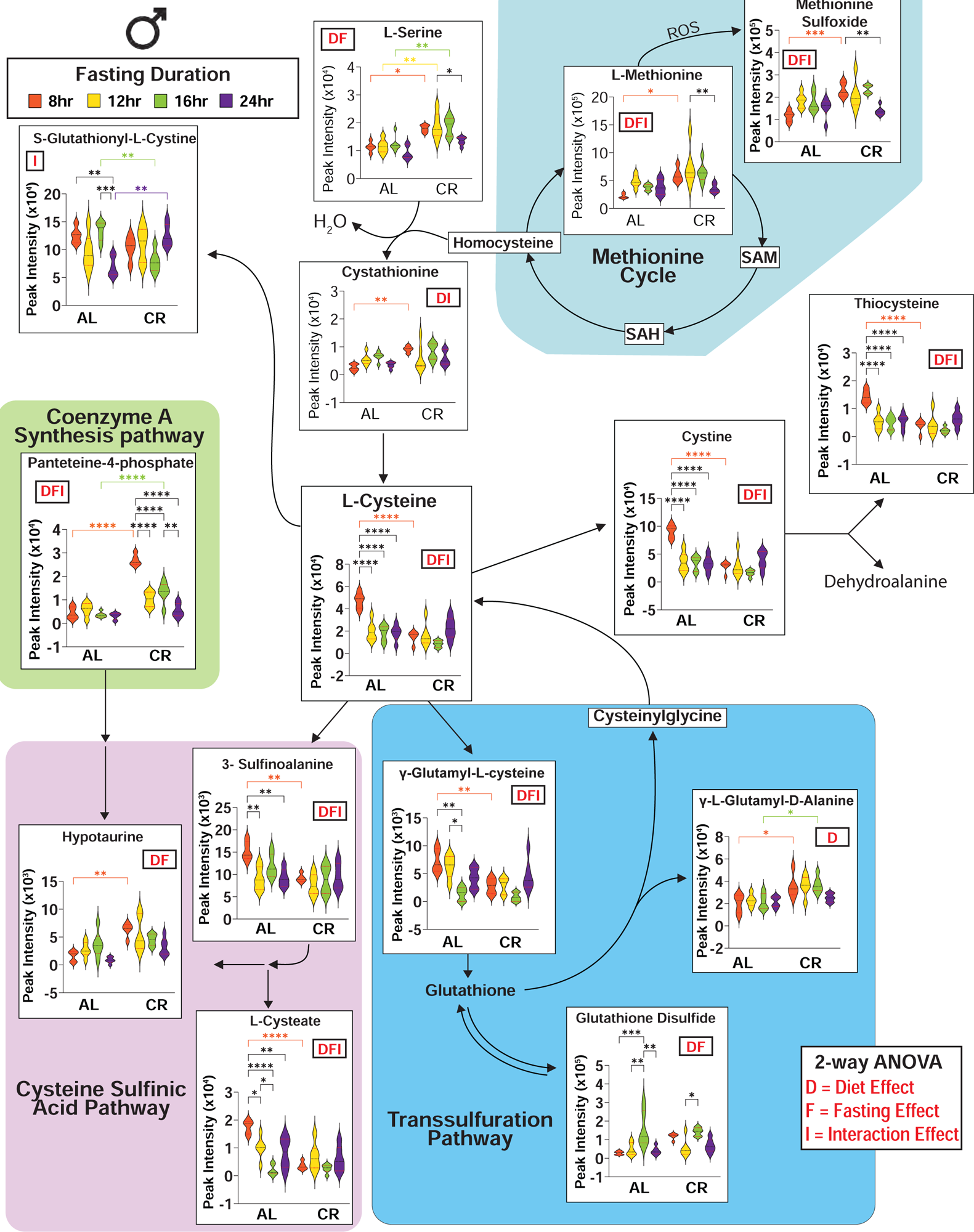
Plasma metabolite data involved in cysteine and methionine metabolism from the untargeted metabolomics analysis, related to Figure 4. *p<0.05 Tukey’s test post two-way ANOVA; D/F/I indicate a significant main effect of diet (D), fasting duration (F), or the diet and fasting duration interaction from the two-way ANOVA(I). The overlaid violin plot shows center as median (black lines) and 25^th^-75^th^ percentiles (red lines).

**Supplementary Figure 14.**
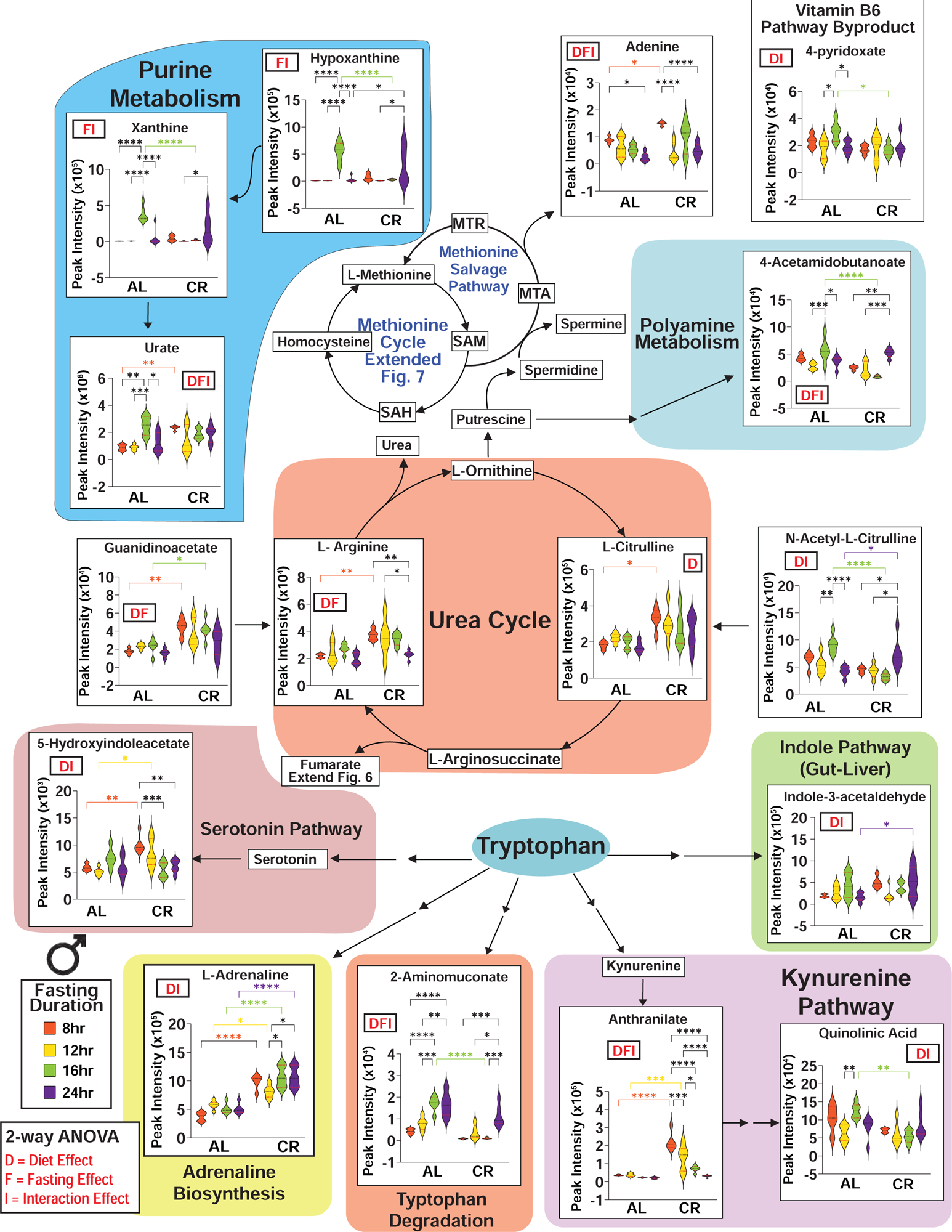
Plasma metabolite data involved in arginine, tryptophan and purine metabolism from the untargeted metabolomics analysis, related to Figure 4. *p<0.05 Tukey’s test post two-way ANOVA; D/F/I indicate a significant main effect of diet (D), fasting duration (F), or the diet and fasting duration interaction from the two-way ANOVA(I). The overlaid violin plot shows center as median (black lines) and 25^th^-75^th^ percentiles (red lines).

**Supplementary Figure 15.**
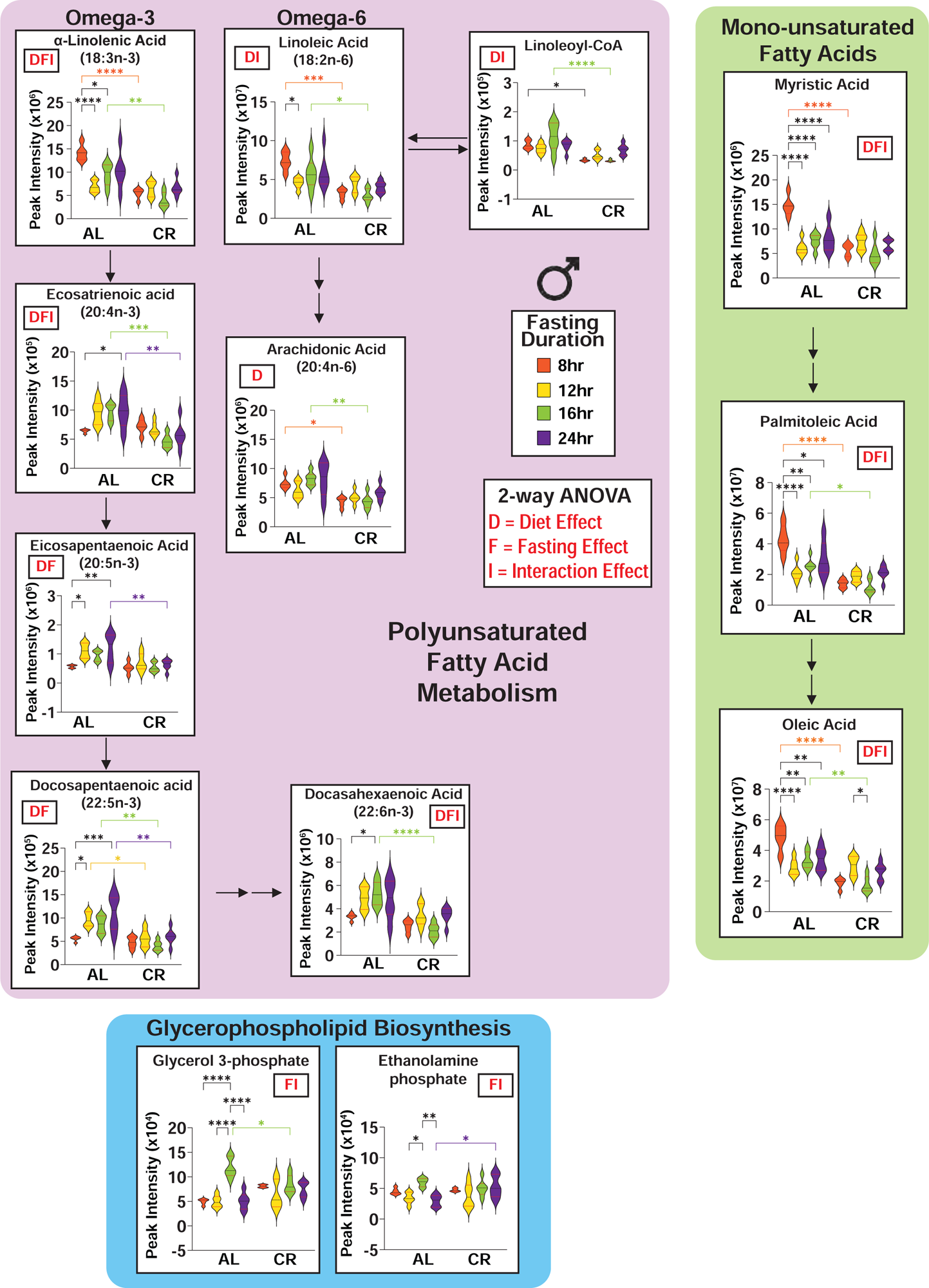
Plasma metabolite data involved in MUFA and PUFA metabolism from the untargeted metabolomics analysis, related to Figure 4. *p<0.05 Tukey’s test post two-way ANOVA; D/F/I indicate a significant main effect of diet (D), fasting duration (F), or the diet and fasting duration interaction from the two-way ANOVA(I). The overlaid violin plot shows center as median (black lines) and 25^th^-75^th^ percentiles (red lines).

**Supplementary Figure 16.**
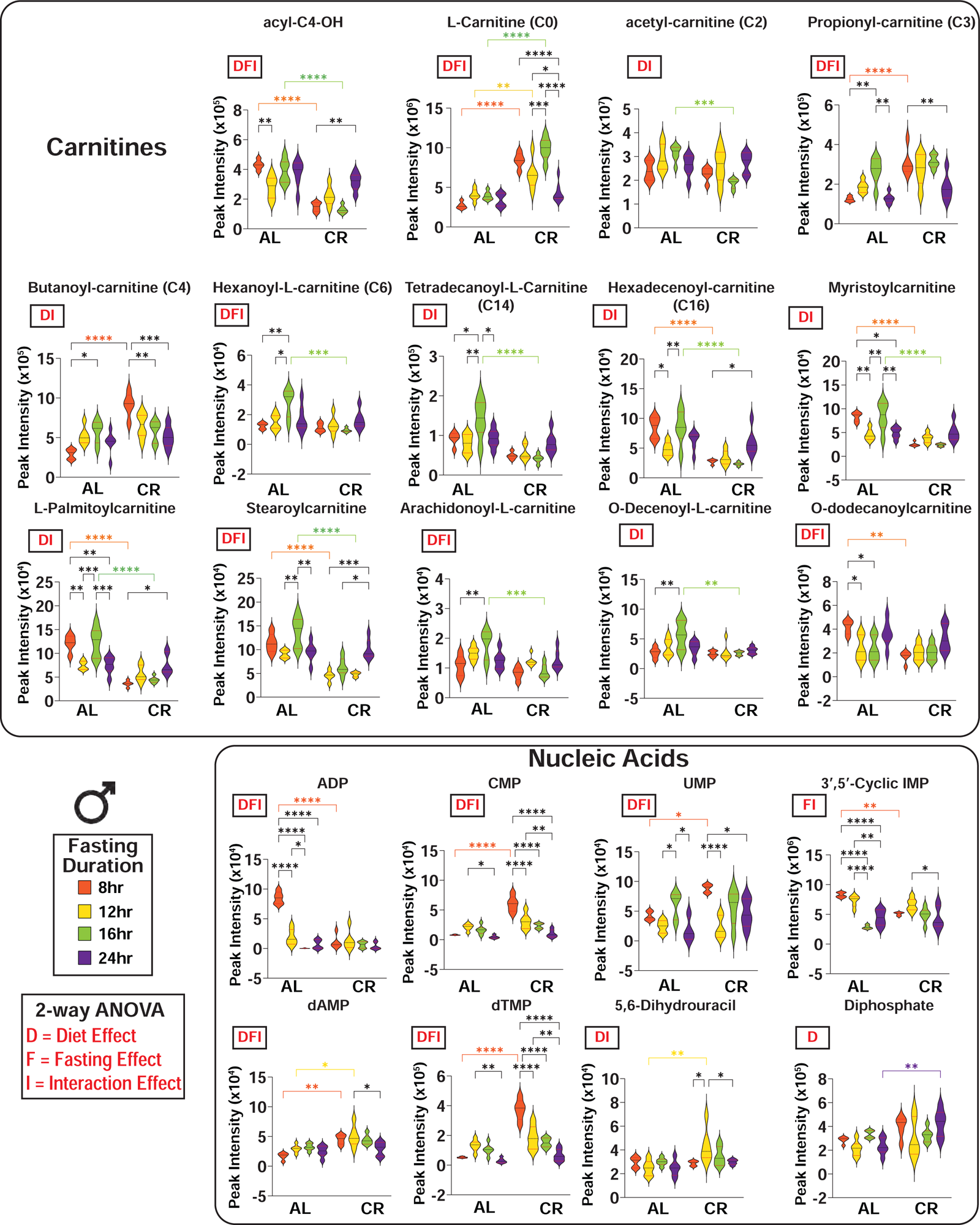
Plasma metabolite data involved in carnitine and nucleic acid metabolism from the untargeted metabolomics analysis, related to Figure 4. *p<0.05 Tukey’s test post two-way ANOVA; D/F/I indicate a significant main effect of diet (D), fasting duration (F), or the diet and fasting duration interaction from the two-way ANOVA(I). The overlaid violin plot shows center as median (black lines) and 25^th^-75^th^ percentiles (red lines).

**Supplementary Figure 17.**
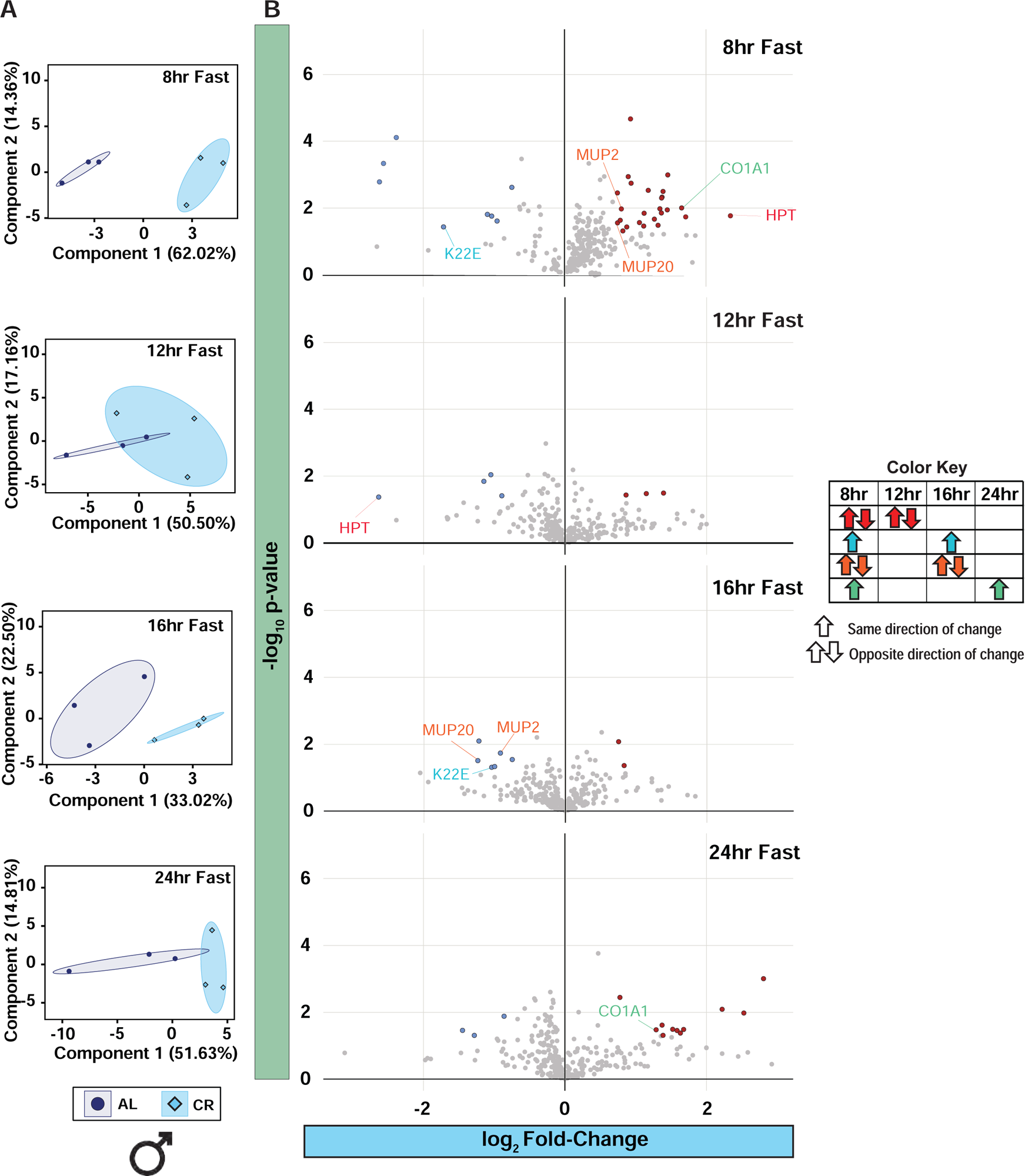
Calorie restriction has minimal effects on the plasma proteome, related to Figure 4. (**A**) Untargeted proteomics were performed on the plasma of male C57BL/6J mice fed AL and CR diet from the Morning Study (n = 5-6 biologically independent mice per diet). PCA of plasma proteins with AL and CR mice collected after an 8hr, 12hr, 16hr and 24hr fast. (**B**) Volcano plot of significantly increased plasma proteins. Volcano plots show the statistical significance (p-value; y-axis) versus magnitude of change (fold-change (log_2_; x-axis). Significantly decreased metabolites are colored blue and significantly increased metabolites are colored red. Metabolite names are colored coded based on significant change between AL and CR at more than one time point (color key).

**Supplementary Figure 18.**
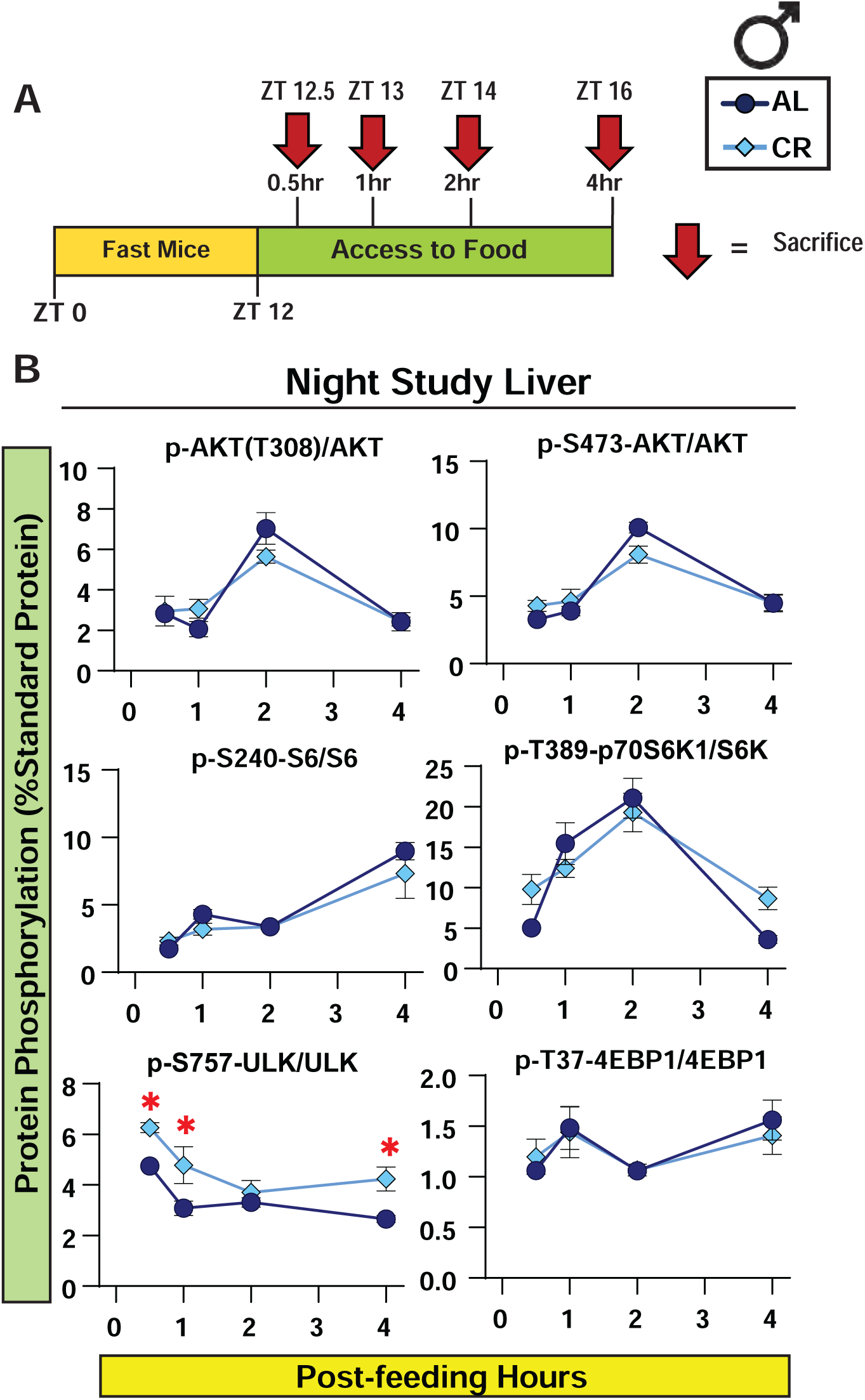
Liver mTORC1 activity is dependent on feeding status, related to Figure 5. (A-B) Experimental design. Food was removed from Night-Fed mice at ZT 0 and all mice were fed at ZT12 which then tissues were collected either at 0.5hr, 1hr, 2hr or 4hr after the first bite of food (B) Liver protein lysate from Night-Fed was analyzed by Western blotting and quantified using ImageJ; y-axis reflects normalization of each phosphoresidue or protein to a liver standard run in duplicate on every gel (n=5 AL-fed and CR-fed biologically independent mice per time point; *p<0.05, Sidak’s test post 2-way ANOVA). Data represented as mean ± SEM.

**Supplementary Figure 19.**
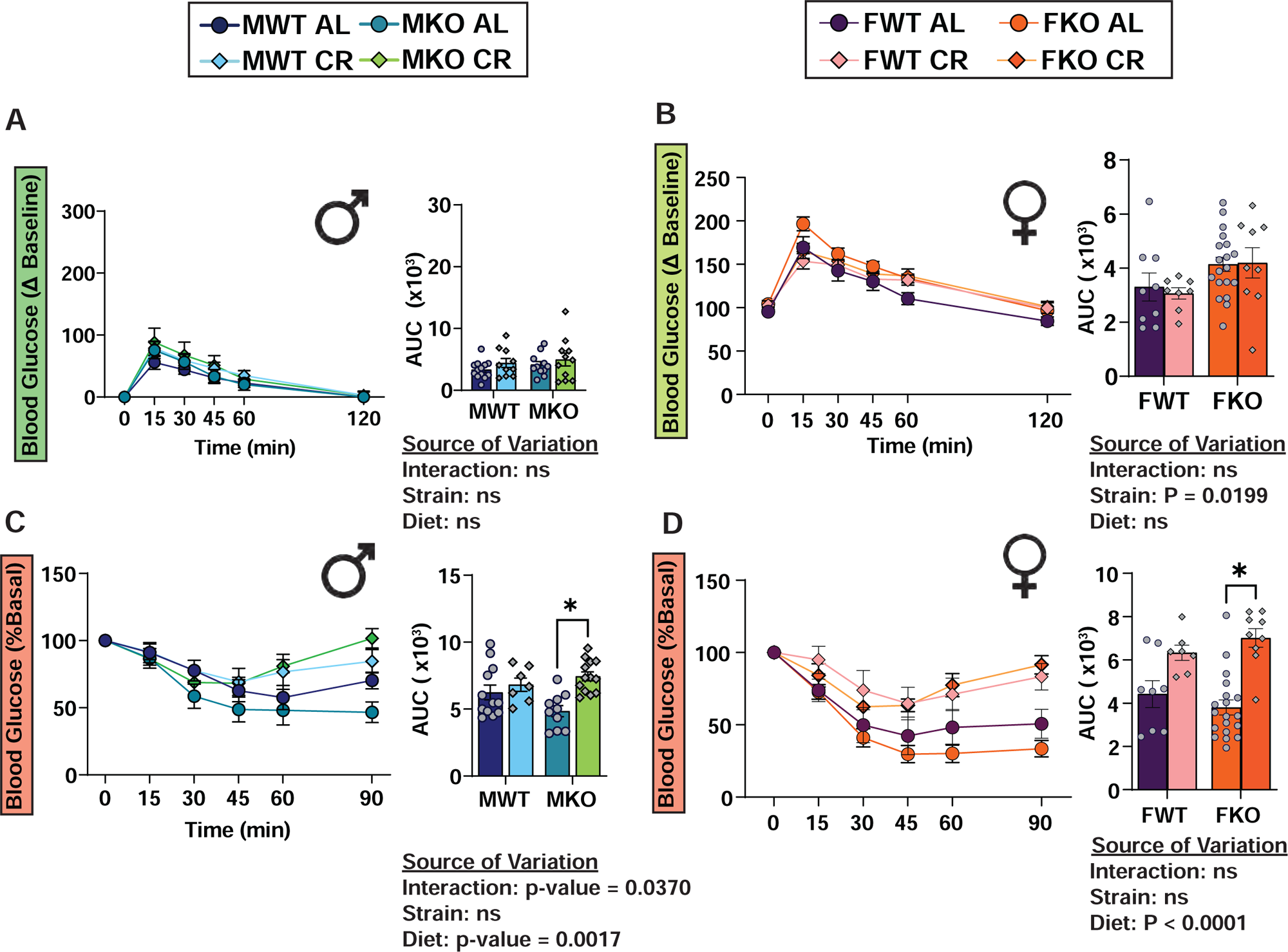
Glucose and insulin tolerance test in Tsc1 (KO) after an 8hr fast, related to Figure 6. (**A-B**) Glucose tolerance tests in male (**A**) and female (**B**) mice lacking hepatic *Tsc1* (KO) and their wild-type (WT) littermates. (**C-D**) Insulin tolerance tests in male (**C**) and female (**D**) mice lacking hepatic *Tsc1* (KO) and their wild-type (WT) littermates. (A,C) MWT AL, n = 13; MWT CR, n = 10; MKO AL, n = 11; MKO CR, n = 10 biologically independent mice. (B,D) FWT AL, n = 9; FWT CR, n = 8; FKO AL, n = 20; FKO CR, n = 9 biologically independent mice. (A-D) statistics for the overall effects of genotype, diet, and the interaction represent the p value from a two-way ANOVA; *p<0.05, from a Sidak’s post-test examining the effect of parameters identified as significant in the 2-way ANOVA. Data represented as mean ± SEM.

## SUPPLEMENTAL TABLE LEGENDS

**Supplementary Table 1. Complete dataset containing raw plasma metabolite values identified by untargeted metabolomics, related to Figure 4**.

**Supplementary Table 2. Cleaned and normalized dataset generated from the raw metabolomics dataset (Supplementary Table 1) containing plasma metabolites identified by untargeted metabolomics. This data was treated as described in the methods section and used in all statistical analyses.**

**Supplementary Table 3. Complete dataset containing raw plasma protein values identified by untargeted proteomics.**

**Supplementary Table 4. Cleaned, filtered, and normalized dataset generated from the raw proteomics dataset (Supplementary Table 2) containing plasma proteins identified by untargeted proteomics. This data was treated as described in the methods section and used in all statistical analyses.**

